# Ototoxicity-induced inner-hair-cell specific dysfunction degrades neurometric modulation detection in noise without altering peripheral tuning

**DOI:** 10.64898/2026.08.20.746057

**Authors:** David Axe, Vijaya Prakash Krishnan Muthaiah, Afagh Farhadi, Michael G. Heinz

**Author notes:** Present address: MathWorks, Natick, MA 01760, USA.

## Abstract

Sensorineural hearing loss can result from different pathologies, but the primary diagnostic method is a threshold-based audiogram, which is insensitive to some forms of cochlear dysfunction. Individuals may experience difficulty understanding speech in noise despite normal audiometric thresholds. Because most cochlear insults damage both inner (IHCs) and outer hair cells (OHCs), the contribution of IHC dysfunction to auditory-nerve coding has been difficult to isolate. We used the IHC-selective ototoxicity of carboplatin in chinchillas to examine how IHC dysfunction, with preserved OHC function, affects temporal-envelope coding in auditory-nerve fibers (ANFs). Carboplatin produced 10–20% IHC loss with stereocilia damage in surviving IHCs, while OHC-dependent measures such as DPOAEs and ANF thresholds were unchanged. Suprathreshold ABR wave 1 was reduced, whereas wave 5 was preserved, suggesting central compensation. Both spontaneous and driven firing rates decreased following exposure. Mean vector strength to amplitude-modulated tones was unchanged, but response variability increased. Neurometric analysis (d′) and mutual information showed degraded AM detection in carboplatin-exposed fibers, an effect accounted for by reduced driven rate (i.e., normalizing spike counts across groups removed the group difference). Background noise degraded AM coding similarly in both groups. Pooled-neurometric modeling showed that population redundancy compensated for impaired fibers in quiet, but not in noise, where carboplatin-exposed pools remained worse. These findings indicate that IHC dysfunction degrades envelope coding by reducing neural output rather than by altering temporal synchrony. This study suggests IHC dysfunction is a phenotype consistent with “hidden hearing loss” (but distinct from cochlear synaptopathy), and motivates suprathreshold clinical assays.

## 1- Introduction

Sensorineural hearing loss (SNHL) is the most commonly diagnosed type of permanent hearing loss. It can arise from a variety of insults including noise overexposure, aging, and exposure to ototoxic agents. Clinically, the audiogram is the gold-standard measure for diagnosing hearing loss and is the first indication of SNHL. However, its limitations in assessing suprathreshold auditory function are well established (Liberman et al., 2016). Two patients with identical audiograms might report dramatically different speech intelligibility even after being fitted with hearing aids (Mangold & Leijon, 1979; Crandell, 1991; Smoorenburg, 1992; Engler et al., 2024; Smith et al., 2024; Lutfi et al., 2026). Similarly, a common complaint amongst patients diagnosed as “normal hearing” is poor intelligibility of speech in noisy environments 8/20/2026 4:52:00 PM. This patient-to-patient variability suggests differences in the underlying mechanisms of SNHL that need to be investigated further.

The complex structure of the cochlea provides many different locations where damage or disruption of various processes can result in impaired hearing ability. Specifically, the mechano-sensory hair-cells are particularly susceptible to damage and do not naturally regenerate in mammals (Clark, 1991; Kiang et al., 1976; Liberman & Beil, 1979; Liberman & Kiang, 1978). The two varieties of auditory hair-cells each serve a unique role in normal-hearing function, and damage or loss of hair-cells is a common cause of SNHL. Outer hair-cells (OHCs) are mechanically responsive to sound stimulation, and their cell body lengthens or contracts in response to changes in membrane voltage (Zheng et al., 2000). This mechanical action contributes additional energy to local basilar-membrane vibrations and improves both cochlear sensitivity and frequency selectivity. Inner hair-cells (IHCs) transduce the incoming pressure wave into neural output along the auditory nerve. Each IHC is innervated by 10-30 spiral ganglion neurons (SGNs) (Liberman et al., 1990). SGNs are bipolar and receive input from only one IHC. These neurons project type-I auditory-nerve fibers (ANFs), which comprise ∼90% of the afferent fibers. The remaining ∼10% are type-II ANFs, which each innervate multiple OHCs, have small unmyelinated axons, and are not known to be sound responsive (Spoendlin, 1969). Type-I ANFs represent the sole source of auditory information to the brain, and any information necessary for normal auditory function must be encoded in their responses. This makes the nerve an ideal site for investigating functional changes in the periphery. Any significant alteration of peripheral function will cause changes in neural coding at the level of the nerve, which will in turn affect the function of processing centers further along the auditory pathway.

Under normal-hearing conditions, ANFs are sharply tuned and most sensitive to a particular frequency (defined as characteristic frequency, CF), which corresponds to the location of innervation along the basilar membrane. Driven spikes of ANFs will synchronize to temporal features of sound stimuli particularly for periodic stimuli; in particular, spike timing will correspond to a particular phase of the stimulus, termed phase-locking. In chinchillas, ANFs will phase-lock to frequencies of up to 2 kHz (Verschooten et al., 2019). ANFs will phase-lock to both the temporal envelope and fine-structure of a stimulus, and proper coding of both features is important for normal auditory function with modulation coding predicted to be important for speech coding in noise (Swaminathan & Heinz, 2012; Verschooten et al., 2019).

In cases of SNHL where OHCs are damaged or destroyed, the normally narrow tuning of ANFs is broadened, and thresholds become elevated. Broadened tuning has been shown to “enhance” envelope coding by increasing the strength of phase-locking (Kale & Heinz, 2010, 2012), but in the presence of background noise fibers impaired in this way show poorer modulation detection (Sayles et al., 2014) and poorer phase-locking to pure-tone carrier frequency (Henry & Heinz, 2012). In cases where IHC function is impaired, the spontaneous and driven spike output of ANFs are reduced, and the slopes of their input-output functions are shallower (Liberman & Dodds, 1984; Liberman & Kiang, 1984) Because many of the pathologies that disrupt the function of one type of hair-cell will affect the other, it is hard to investigate the specific effects of IHC disruption on temporal coding in the nerve without the confound of degraded tuning due to reduced OHC function.

Carboplatin is a platinum-based chemotherapy drug commonly prescribed for ovarian and other cancers with well-known ototoxic side effects (Qaddoumi et al., 2012). Similar to other platinum-based drugs, carboplatin’s cytotoxicity is believed to stem from the cross-linking of DNA and inhibiting DNA repair and replication. In chinchillas, the ototoxicity of the drug is specific to IHCs (Brock et al., 2012; Henderson et al., 1999). In chinchillas, the ototoxicity of the drug is specific to IHCs (Wake et al. 1994; Trautwein et al. 1996; Ding et al. 1999; El-Badry and McFadden 2009; Wang et al. 1997; Lobarinas et al. 2013, 2015). At high doses, OHCs may also be damaged or destroyed, but typically this occurs after complete loss of IHCs (D. L. Ding et al., 1999). The extent of IHC loss is dosage dependent and affects IHCs along the entire length of the cochlea (D. L. Ding et al., 1999; Trautwein et al., 1996). Remaining IHCs are also affected, showing disrupted stereocilia bundles (Wake et al., 1994). The amplitude of evoked potentials is reduced after exposure, but thresholds are insensitive (El-Badry & McFadden, 2009; Trautwein et al., 1996). Behavioral studies have shown similar results. Even in cases where 80% loss of IHCs was reported but neural correlates were not known, behavioral pure-tone detection thresholds were not significantly elevated (Lobarinas et al., 2013, 2015).

The purpose of the present study is to utilize carboplatin’s IHC-specific ototoxicity as a means to investigate how disruption of IHC function, while preserving normal OHC function, affects the coding of temporal envelope features in the auditory nerve in quiet and in the presence of background noise (the condition in which listeners with SNHL struggle the most) and to use signal detection theory to evaluate the discriminability of these signals.

## 2- Methods

### 2-1- Animal subjects

Subjects were adult male chinchillas (400-700 g, 23 control, 23 carboplatin (CA) exposed). Animals were 6 to 12 months of age at the time of carboplatin exposure. All animals were screened for normal hearing prior to inclusion in the study using the evoked potential protocol outlined below. All procedures were approved by the Purdue University IACUC committee.

### 2-2- Carboplatin (CA) exposure

Following baseline measurements of hearing ability, animals received a single injection of carboplatin (38 mg/kg, IP). Previous studies in chinchillas have shown that this dosage induces an average loss of ∼20% of IHCs along the entire length of the cochlea and disruption of stereocilia in surviving IHCs (Trautwein et al., 1996). Following injection, subjects were returned to their housing facility and allowed to recover for at least 2 weeks before an acute neurophysiological procedure was performed.

### 2-3- Non-invasive assays of hearing status: auditory brainstem responses (ABRs) and distortion product otoacoustic emissions (DPOAEs)

Procedures for non-invasive assays were conducted using techniques defined in previous studies (Henry et al., 2011; Zhong et al., 2014). Animals were kept under anesthesia during all physiological recordings. Anesthesia was induced using ketamine (40 mg/kg, IP) and xylazine (4 mg/kg, SQ). Heart rate and blood oxygen level were monitored with a pulse-oximeter, and an internal body temperature of 37° C was maintained via an electric heat-blanket controlled via feedback from an internal thermometer (Harvard Apparatus). Physiological saline and lactated Ringer’s solution were given regularly during anesthetized procedures to prevent dehydration (∼1 ml/h). All physiological recordings were conducted in an electrically shielded, double-walled sound-attenuating booth.

Prior to carboplatin exposure, baseline physiological measures of hearing were collected to ensure normal hearing at entry into the study using DPOAEs and ABRs. Basic hearing ability was again assessed in the same manner just prior to single-unit recordings. For these procedures, sound stimuli were presented through two sealed transducers inserted into the ear canal (Etymotic ER-2, Elk Grove Village, IL, USA). Stimuli were calibrated, and OAE responses were recorded using an integral microphone (Etymotic ER-10B, Elk Grove Village, IL, USA). DPOAE stimuli consisted of two tones, f1 and f2, where f2>f1, and with a frequency ratio of 1.21. DPOAEs were recorded across a range of frequencies with f2s from .5 to 10 kHz. Both tones were presented at 75 dB SPL, and DPOAE amplitudes were measured at the frequency of the strongest distortion product (2f1-f2).

ABRs stimuli were presented through the same speaker system used for DPOAEs. Stimuli consisted of 10-ms tone pips presented at .5, 1, 2, 4, and 8 kHz. Responses to each frequency were recorded at sound levels starting at 80 dB SPL and decreasing in 10 dB steps to 10 dB below the visual threshold. Responses were recorded through needle electrodes inserted into the scalp at the midline between the dorsal bulla (non-inverting), posterior to the pinna of the ear being recorded (inverting), and at the bridge of the nose (ground). Individual responses were amplified by x20,000 (World Precision Instruments ISO-80, Sarasota, FL; Dagan 2400A, Minneapolis, MN), band-pass filtered from 0.3 to 3 kHz (Krohn-Hite 3550, Brockton, MA), and digitally sampled at a rate of 48828.125 Hz (TDT RP2.1, Alachua, FL). A set of 500 responses of each polarity were averaged to obtain the response at a given stimulus frequency and sound level.

### 2-4- Surgical preparation and single-unit recording

Procedures for surgical preparation and single-unit isolation were conducted using techniques defined in previous studies (Kale and Heinz, 2010; Henry et al., 2016 – JN DT paper; Parida and Heinz, 2022 – JN paper). During acute procedures, following DPOAEs and ABRs, anesthesia was maintained for the remainder of the preparation with pentobarbital (∼7.5 mg/kg/hr, IV). In these procedures, a tracheotomy was performed to allow for a low-impedance airway. The skin and muscles overlying the skull were reflected to expose the ear canals and bulla. The bulla was vented with a 30 cm long polyethylene tube to maintain the middle ear pressure. Hollow ear bars were placed in the ear canal as part of the stereotaxic apparatus holding the animal in place and coupled with the tympanic membrane in order to form a closed-field system. A craniotomy was made in the posterior fossa, exposing the cerebellum, which was then partially aspirated to reveal the cochlear nucleus just dorsal to the internal auditory meatus. The brainstem was gently retracted medially with cotton pellets to reveal the AN as it exits the internal auditory meatus. Recordings from individual ANFs were made using glass 10-30 MΩ micropipette electrodes filled with 3 M NaCl. Electrodes were placed under visual control as close to the opening of the auditory meatus as possible and then advanced into the nerve using a hydraulic Micropositioner (Kopf). The electrode signal was amplified (Dagan 2400A, Minneapolis, MN) and bandpass filtered from .03 to 6 kHz (Krohn-Hite 3550, Brockton, MA) before spike timing information was recorded. Spike timing was measured with 10-μs resolution using a time-amplitude window discriminator (Bak Electronics, Mount Airy, MD, USA). A second low-impedance silver electrode was placed on the round window via a hole opened in the caudal portion of the bulla. This electrode was used throughout the experiment to monitor compound action potentials as a means of assessing the ongoing health of the periphery during the procedure. If CAP thresholds elevated more than 10 dB above the baseline, the experiment was ended.

Stimuli were delivered through dynamic speakers (DT-48, Beyer Dynamic, Farmingdale, NY, USA) coupled to the tympanic membrane via the hollow ear bars. The acoustic system was calibrated for each preparation just prior to recording using a probe-tube microphone (Etymotic ER-7C, Elk Grove Village, IL, USA) placed adjacent to the tympanic membrane. Simultaneous generation of acoustic stimuli and data collection was controlled using custom software running in MATLAB (The Mathworks, Natick, MA, USA) that was integrated with commercial hardware (Tucker-Davis Technologies, Alachua, FL, USA; National Instruments, Austin, TX, USA).

Individual ANFs were isolated by advancing the electrode slowly through the nerve while presenting a broadband noise probe stimulus. Once isolated, ANFs were characterized using a standardized protocol. Threshold and CF were measured using an automated tracking algorithm to generate a tuning curve (Chintanpalli & Heinz, 2007). This algorithm determined the minimum sound level for a 50-ms tone to elicit at least one more spike than a subsequent silence. Fiber CF, threshold, and Q_10_ (ratio of CF to tuning-curve bandwidth 10 dB above threshold) were estimated from the tuning curve. Fiber spontaneous rate (SR) was then measured by recording spikes during a 20-second period of silence. CF-tone rate-level functions were then collected. Responses to 50-ms CF tones were measured with sound levels ranging from ∼0 dB SPL until driven rate (DR) saturation was visually identified. ANFs were further characterized by collecting CF-tone post-stimulus-time (PST) histograms. PST histograms were constructed from responses to 300 repetition of a 50 ms CF-tone.

### 2-4- Amplitude modulation coding

Following general characterization, amplitude modulation (AM) coding was measured using sinusoidally amplitude-modulated (SAM) tones. Since the strength of AM phase-locking is heavily dependent on sound level, the best-modulation level (BML) was determined by collecting a rate-level curve for SAM tones. Stimuli for this SAM tone rate-level function were 600 ms fully modulated tones (modulation depth, m=1) with a 400 ms off time between presentations. SAM tones were presented with a carrier frequency (fc) equal to the CF of the ANF, and the modulation frequency (f_m_) was 50 Hz. Stimuli were presented at sound levels ranging from 5 dB below threshold to 30-40 dB above in 5 dB increments. Stimuli at all levels were repeated 20-30 times. BML was determined to be the sound level with the greatest synchrony to f_m_. AM depth coding was assessed using SAM tones identical to those used for the SAM RL above (fc=ANF’s CF, f_m_ = 50 Hz) but with varying modulation depths (m=[0, .0313, .0625, .125, .25, .5, 1] or m=[-INF, -30, -24, -18, -12, -6, 0] dB). Stimuli were repeated in the presence of broadband noise presented at 10, 15, and 20 dB relative to the tone signal; overall sound level was set to the BML of the fiber. The full set of stimulus parameters was repeated 10-20 times.

### 2-5- Single-Unit Data Analysis

Basic quantification of synchrony of phase-locking was calculated using vector strength (Goldberg & Brown, 1969), also known as the synchronization coefficient, calculated by

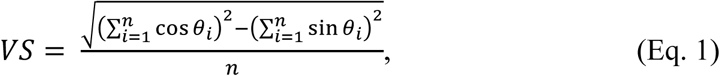

where VS is vector strength, n is the number of spikes across all trials, and θ_i_ is the phase of each spike in radians. One of the main limitations of the VS metric is that it is inaccurate when low numbers of spikes are available and can overestimate synchrony in these conditions. Phase-projected vector strength is a modified form of VS that aims to correct for this by adding in a weighting factor which scales the calculated VS value based on its consistency across multiple repetitions of the same stimulus (Yin et al., 2011). The formula for VS_pp_ is calculated by

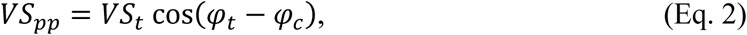

where VS_pp_ is the phase-projected vector strength, VS_t_ is the vector strength per trial, and φ_t_ and φ_c_ are the trial-by-trial and mean phase angle, respectively, in radians, calculated for each stimulus condition, where

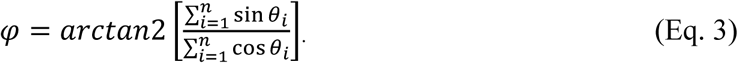

This weighting factor compensates for artifactually high VS values when few spikes are included in the analysis and allows for accurate measures of spike synchrony to be calculated for small sets of spikes even for the response to an individual repetition of the stimulus.

In order to quantify the amount of information that the measured VS_pp_ conveyed about the depth of modulation in the stimulus the metric mutual information (MI) was calculated (Cover & Thomas, 2001). MI is a measure of the mutual dependence between two random variables and quantifies the “amount of information” (in this case calculated as bits) obtained about one random variable when the value of the other is known. The general equation for MI is

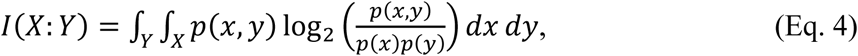

where *p(x,y)* is the joint probability of function of the random variables *X* and *Y*, and *p(x)* and *p(y)* are the marginal probability distribution functions of *X* and *Y*. For the purposes of this study, the variables X and Y represent the measurable properties of modulation depth and VSpp, respectively. The calculated MI quantifies how much information about the modulation depth of the stimulus you gain from knowing the measured VS_pp_.

In order to investigate signal discriminability, neurometric analyses were used (Sayles et al., 2013, 2014). Using signal-detection theory, the discriminability of the response of a modulated tone from an unmodulated tone (modulation detection) and of a modulated tone from a fully modulated tone (modulation discrimination) was quantified as the discrimination index (*d’*):

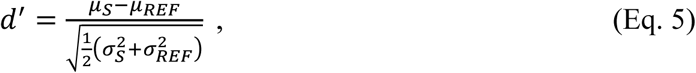

where µ_S_ and µ_REF_ are the mean responses to the stimulus and to the reference condition, respectively, and σ_S_ ^2^ and σ_REF_ ^2^ are their variances (Fig. 1, D). A key assumption of this method is that the distributions being compared are both normal. Due to the stability limitations of single-unit recordings, a limited number of repetitions are possible. In order to generate sufficiently large distributions for neurometric analyses, simulated repetitions were bootstrapped from collected data (Efron & Tibshirani, 1998). All spikes from all repetitions of a given stimulus were pooled, and a number of spikes equal to the mean driven rate of the unit were selected at random without replacement (Fig. 1, A). This produced a single bootstrapped “repetition”. This process was then repeated 5000 times to generate a sufficiently large number of bootstrapped repetitions. VSpp was calculated for each rep, and these values comprised the distributions used in the analyses (Fig. 1, B and C).

**Fig. 1.**
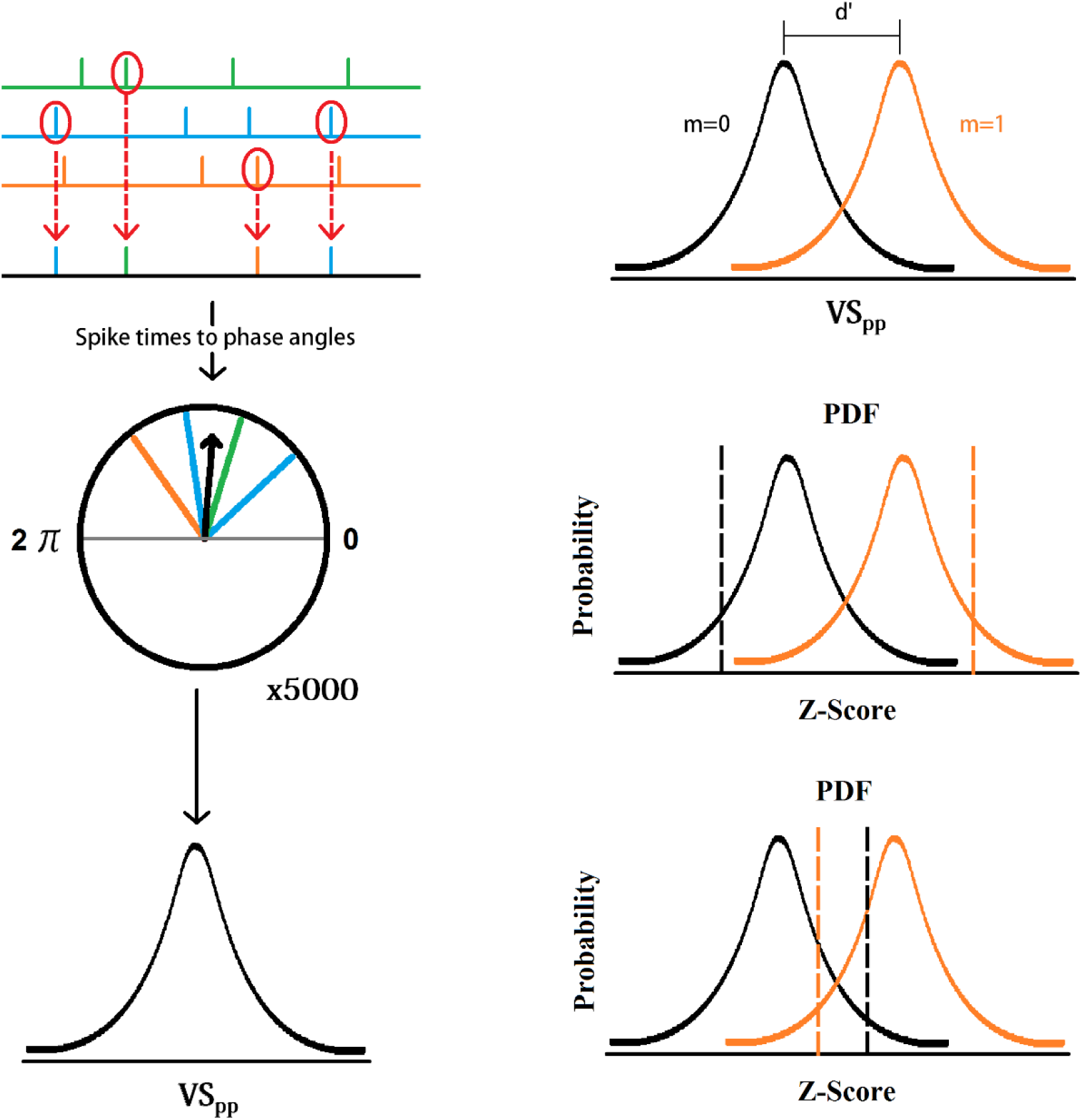
Schematic of neurometric analyses used. A-C (left column), Calculation of VS_pp_ from bootstrapped spike-trains. A, Bootstrapped spike trains are generated by random picking of spikes from measured spike trains (green, blue, and orange). B, The spike times of the bootstrapped spike trains are converted into phase angles relative to the modulation frequency, and the vector average is calculated (VS). C, The bootstrapping process is repeated 5000x, and the calculated phase-projected vector strength (VS_pp_) averages are weighted based on consistency across reps in order to generate the distributions used in the neurometric analysis. **D, discrimination index (*d’*) is calculated by comparing the mean and variance of the modulated probe stimulus being detected (orange) to the mean and variance of the unmodulated reference probe stimulus (black).** *d’* is the difference in the means of the probe and reference divided by the square root of the sum of their variances divided by 2 (Eq. 5). **E and F, Pooled-neurometric analysis is based on comparing the summed responses to a modulated probe stimulus to an unmodulated reference for a pooled group of neurons.** The mean synchrony (VS_pp_) of each neuron in a pool is converted to a Z-score and randomized by the addition of a random number with mean zero and variance equal to the variance of the fiber’s calculated VS_pp_ across all bootstrapped reps. The sum of a pool’s randomized Z-scores can be represented as a probability density function (PDF) with a mean equal to the sum of the mean Z-scores for all fibers included in the pool. For each “trial” of the pooled-neurometric task, a summed Z-score value is generated for both the signal (black dashed line) and reference (red dashed line). If the probe value is greater than the reference value the trial is scored correct (panel E). If the value generated for the reference probe is greater than for the probe value the trial is scored incorrect (panel F).

### 2-6- Pooled Neurometric Analysis

In an effort to quantitatively investigate how modulation information is integrated across a population of ANFs, a pooled-neurometric analysis was performed for a modulation-detection task based on an assumed ideal observer following the methods outlined by Rosen et al. (Rosen et al., 2010, 2012). In this analysis, a simulated two-interval two-choice discrimination task was used. Pool sizes were varied between 1 (individual fiber responses), 2, 5, 10, 20, and 50. Pools were generated by randomly selecting fibers without replacement from the total fiber population. For each fiber included in a given pool, the distributions of VSpp for individual bootstrapped repetitions were then converted into Z-scores. The mean Z-scores for each fiber’s response to each stimulus condition were modified by the addition of a random factor meant to simulate neuronal variability. This random factor was weighted such that the final sum, the random number *X,* had a mean and variance equal to the experimental data that it was drawn from, such that:

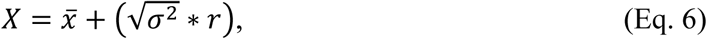

where *x̄ is* the mean Z-score of the VSpp distribution, *σ^2^* is its variance, and *r* is a normally-distributed random number with mean of zero and standard deviation of 1 (generated by MATLAB function *randn*). Randomized responses from each unit within a pool were summed together to form the total pool response. Modulation detection was determined by comparing the pool’s summed response from the modulated stimulus to the summed response to the unmodulated reference. The greater value of the two summed responses was determined to be the pool’s “choice” in the task. In cases where the summed response to the modulated condition was greater than the summed response to the unmodulated reference condition, the pool’s response was taken as correct. For each generated pool, this task was repeated 20 times for each stimulus condition in order to determine that pool’s percent-correct score. This process was repeated 1000 times for the NH and CA-exposed populations with a new randomly generated pool of ANFs each time.

### 2-7- Cochleograms and Scanning Electron Microscopy

Following acute electrophysiological recordings, selected animals were perfused intracardially with normal saline followed by fixation with 10% formalin in phosphate-buffered saline (PBS). Temporal bones were harvested and maintained in fixative at 4°C for 24 h before decalcification in 10% EDTA at 4°C for 4 days. Cochleae were then rinsed in 0.1 M PBS and processed as surface preparations (D. Ding et al., 2001). After staining with Harris hematoxylin, specimens were mounted and examined under a compound microscope (Zeiss Standard). Inner hair cells (IHCs) and each of the three rows of outer hair cells (OHC1–OHC3) were quantified at 0.24-mm intervals along the entire basilar membrane using 400× magnification. Cell counts were analyzed using custom software to generate cochleograms showing the distribution of missing IHCs and OHCs (average across OHC rows) as a function of cochlear position relative to the apex (D. Ding et al., 2013).

Scanning electron microscopy (SEM) was used to evaluate cochlear morphology following carboplatin treatment using methods similar to those described by (Wake et al., 1994). Specimens were fixed in 2.5% glutaraldehyde in 0.1 M sodium cacodylate buffer, post-fixed in 1% osmium tetroxide, dehydrated in a graded ethanol series, and transferred into a Tousimis 931 CPD for critical point drying. Dried specimens were coated with platinum in a Cressington 208HR sputter coater. Electron microscopy and SEM sample preparation was performed using instrumentation in the Purdue Electron Microscopy Center (RRID:SCR_022687).

## 3- Results

### 3-1- Anatomical changes following CA exposure

Carboplatin exposure resulted in selective inner hair cell (IHC) pathology while outer hair cells (OHCs) remained largely intact (Fig. 2). Bright-field microscopy showed missing IHCs despite preserved OHC rows (Fig. 2.A), consistent with the selective vulnerability of IHCs to carboplatin (D. -L. Ding et al., 1999; Hofstetter, Ding, Powers, et al., 1997a; Hofstetter, Ding, & Salvi, 1997; Lobarinas et al., 2013, 2015, 2020; Salvi et al., 2017; Takeno et al., n.d.; Trautwein et al., 1996; Wake et al., 1994; Wang et al., 1997b). The extent of IHC loss varied somewhat across animals, but was generally consistent across frequency region for the mild dose used in the present study with ∼10 to 15% IHC loss across the cochlea (Fig. 2.B).

**Fig. 2.**
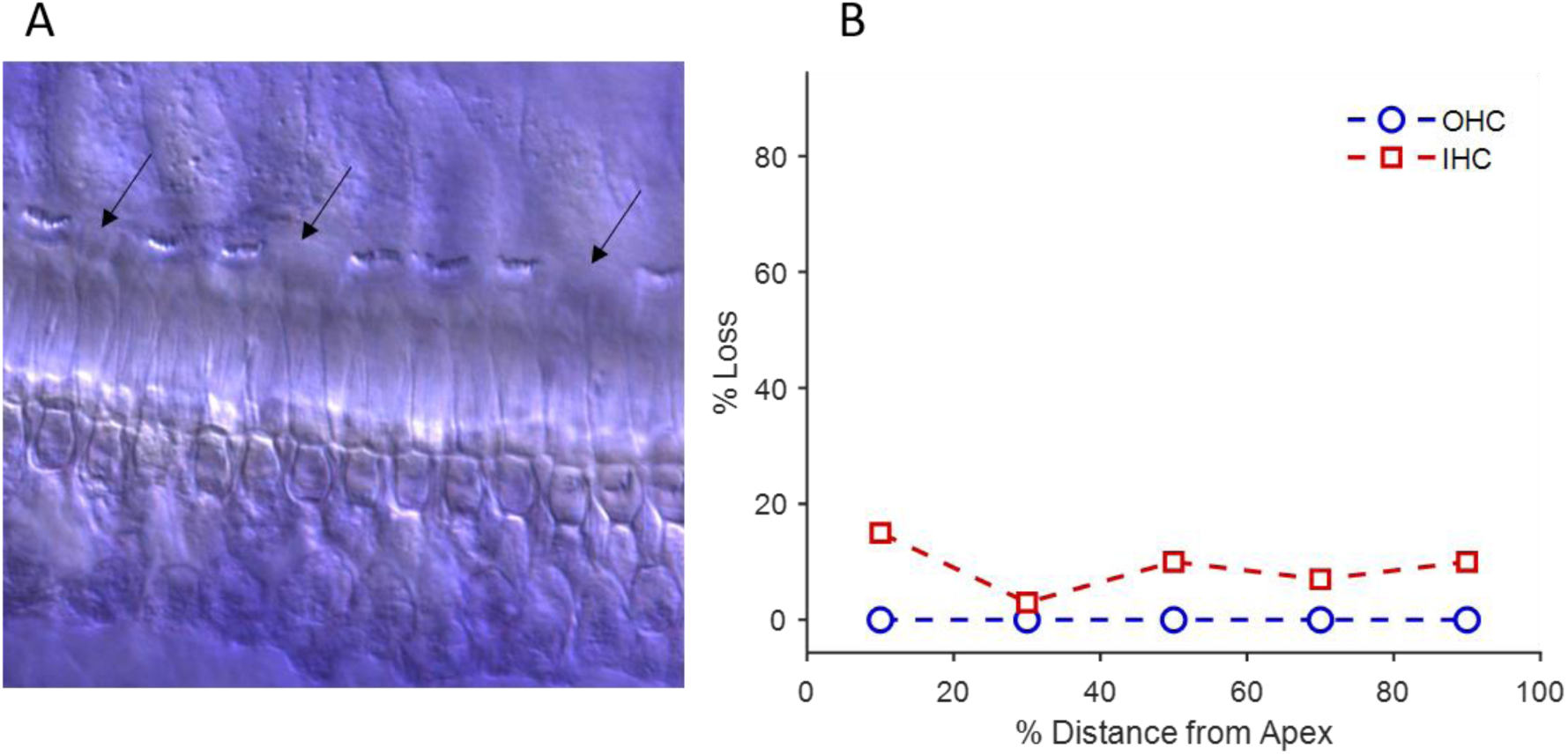
Example of anatomical changes following CA exposure. A) Bright-field microscopy of the organ of Corti. The majority of IHCs remain intact, with black arrows indicating missing IHCs. B) Representative cochleogram of IHC and OHC loss relative to location within the cochlea.

In contrast, OHC loss remained negligible at all cochlear locations examined. Preservation of OHCs in the same tissue argues against the IHC disruption being a fixation or dissection artifact. This findings are in line with previous studies that have counted remaining hair-cells following exposure to identical (Ding et al., 1999; Takeno et al., 1994; Trautwein et al., 1996) or larger dosages of CA (Hofstetter, Ding, Powers, et al., 1997b; Lobarinas et al., 2013, 2015) and showed no or little loss of OHCs.

SEM imaging suggested that cochleograms likely underestimate the extent of IHC pathology (Fig. 3). Although only a modest fraction of IHCs were absent, many of the remaining IHCs showed clear abnormalities of their stereociliary bundles. These abnormalities included partial loss of stereocilia, missing stereocilia, fractured stereocilia, and apparent displacement or uprooting of stereocilia. In contrast, OHC stereociliary bundles appeared largely unaffected. As illustrated in Fig. 3A, OHC stereociliary bundles remained intact and well organized in the same preparation despite clear disruption of the adjacent IHC stereocilia. Thus, this mild carboplatin exposure resulted not only in loss of some IHCs, but also substantial structural damage among surviving IHCs. This selective pattern of IHC damage with relative preservation of OHCs is consistent with previous reports of carboplatin-induced selective IHC ototoxicity in chinchillas (Wake et al., 1994).

**Fig. 3.**
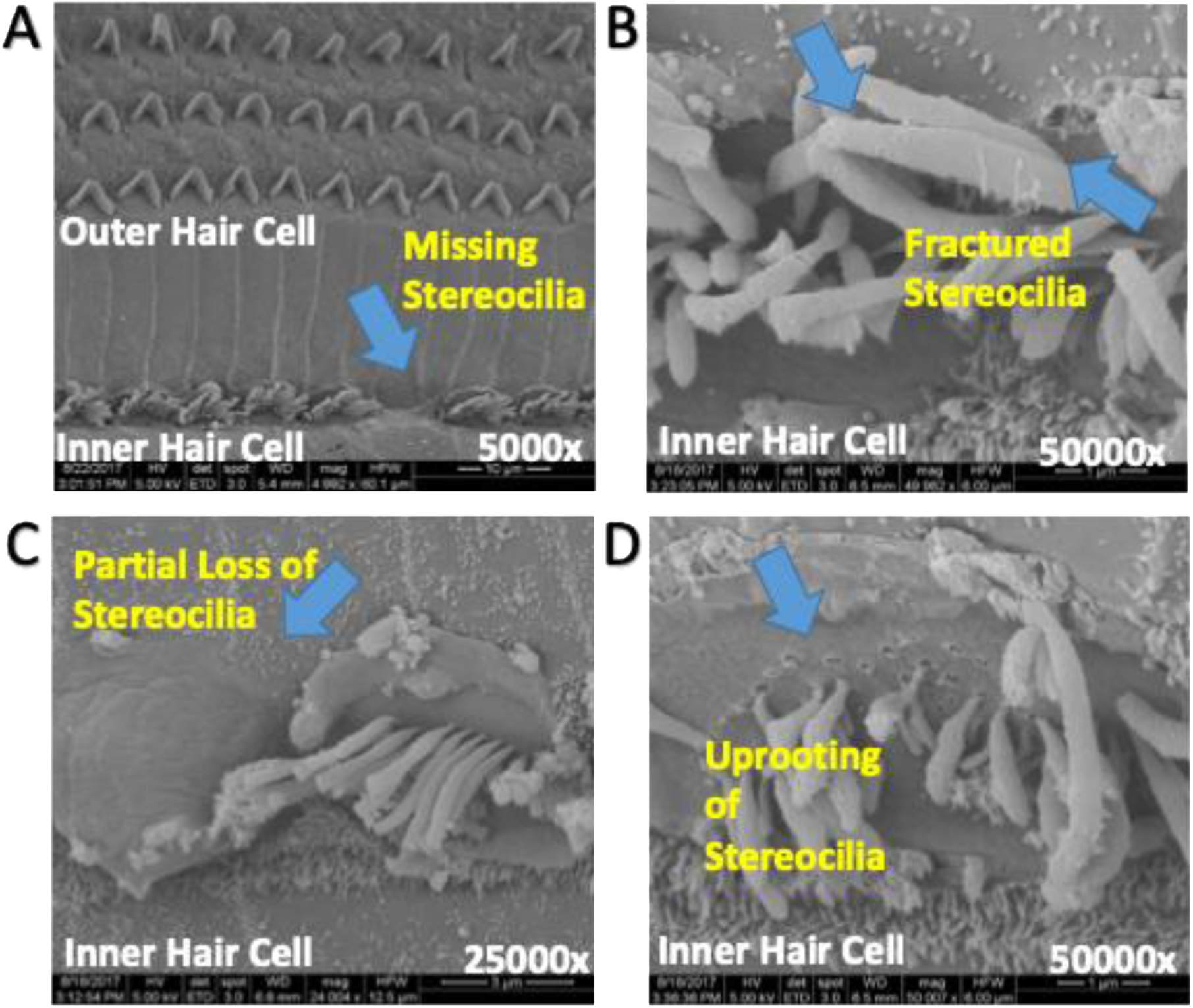
Effect of carboplatin on stereocilia integrity in chinchilla inner hair cells (scanning electron microscopy). (A) Outer hair cell (OHC) stereocilia remain intact and well organized, whereas inner hair cell (IHC) stereocilia show marked disarrangement and missing stereocilia. (B–D) Higher-magnification images of IHCs demonstrate additional stereocilia abnormalities, including fractured stereocilia (B), partial stereocilia loss (C), and uprooting of stereocilia (D). Overall, these images demonstrate substantial IHC stereocilia damage despite preservation of OHC stereocilia following carboplatin exposure.

### 3-2- Noninvasive physiological assessment following carboplatin exposure

Exposure to CA did not affect DPOAE amplitude (Fig. 4, Tukey’s HSD, *Q*=2.35, *P*=0.23). DPOAEs are produced directly by the stimulus-evoked nonlinear mechanical activity of OHCs. As such DPOAE amplitudes have been shown to be sensitive to even mild amounts of SNHL involving OHC dysfunction. The present findings indicate that OHC functionality has not been significantly altered by CA exposure, which is confirmed by histology results cochleogram in section 3-1.

**Fig. 4.**
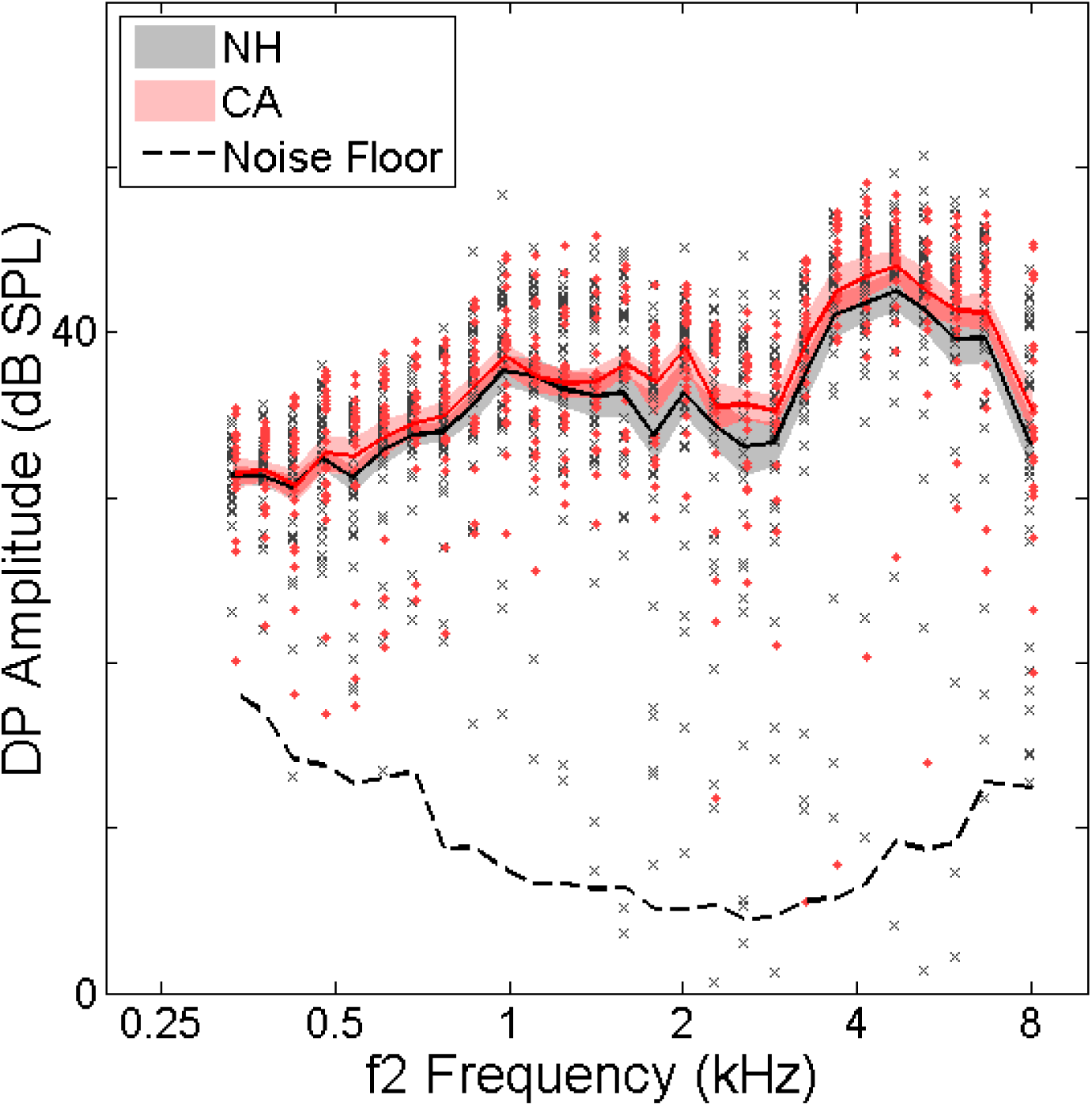
Average DPOAE amplitudes did not change following CA exposure. NH (black) and CA (red) exposed animals showed no difference in DP amplitudes (plotted relative to f2 frequency). Solid lines show mean DP amplitude for each F2 frequency. Shaded regions depict 95% confidence intervals of the mean. Grey x’s and red diamonds show amplitudes for individual NH- and CA-exposed animals, respectively. The Dashed line illustrates the average measured noise floor.

ABR thresholds did not significantly change on average (Fig. 5, Tukey’s HSD, *Q*=2.36, *P*=0.32). Wave-1 of the ABR complex is generated directly by the summed potential of synchronous firing across ANFs (Buchwald & Huang, 1975). Following CA-exposure, wave-1 amplitudes were reduced (Fig. 5 insert). At the highest sound level recorded, 90 dB SPL, the average response across CA-exposed ears was 33% smaller in amplitude compared to the average response in normal ears (NH: 1.98 uV, CA: 1.33 uV, *T*_102_ = 3.46, *P* = .00079). Wave-2, generated in the cochlear nucleus, showed a similar reduction in amplitude following exposure (data not shown). In the standard electrode configuration used during these recordings, wave-3 was often merged either partially or wholly with either wave-2 or wave-4 depending on stimulus frequency, and thus it was not possible to reliably measure wave-3 amplitude. Wave-4 and wave-5, generated in the auditory midbrain, did not show any significant reduction in amplitude following exposure (Data not shown, Tukey’s HSD, *Q*=2.35, *P*=0.92). These findings suggest some form of central compensation occurs following CA-exposure.

**Fig. 5.**
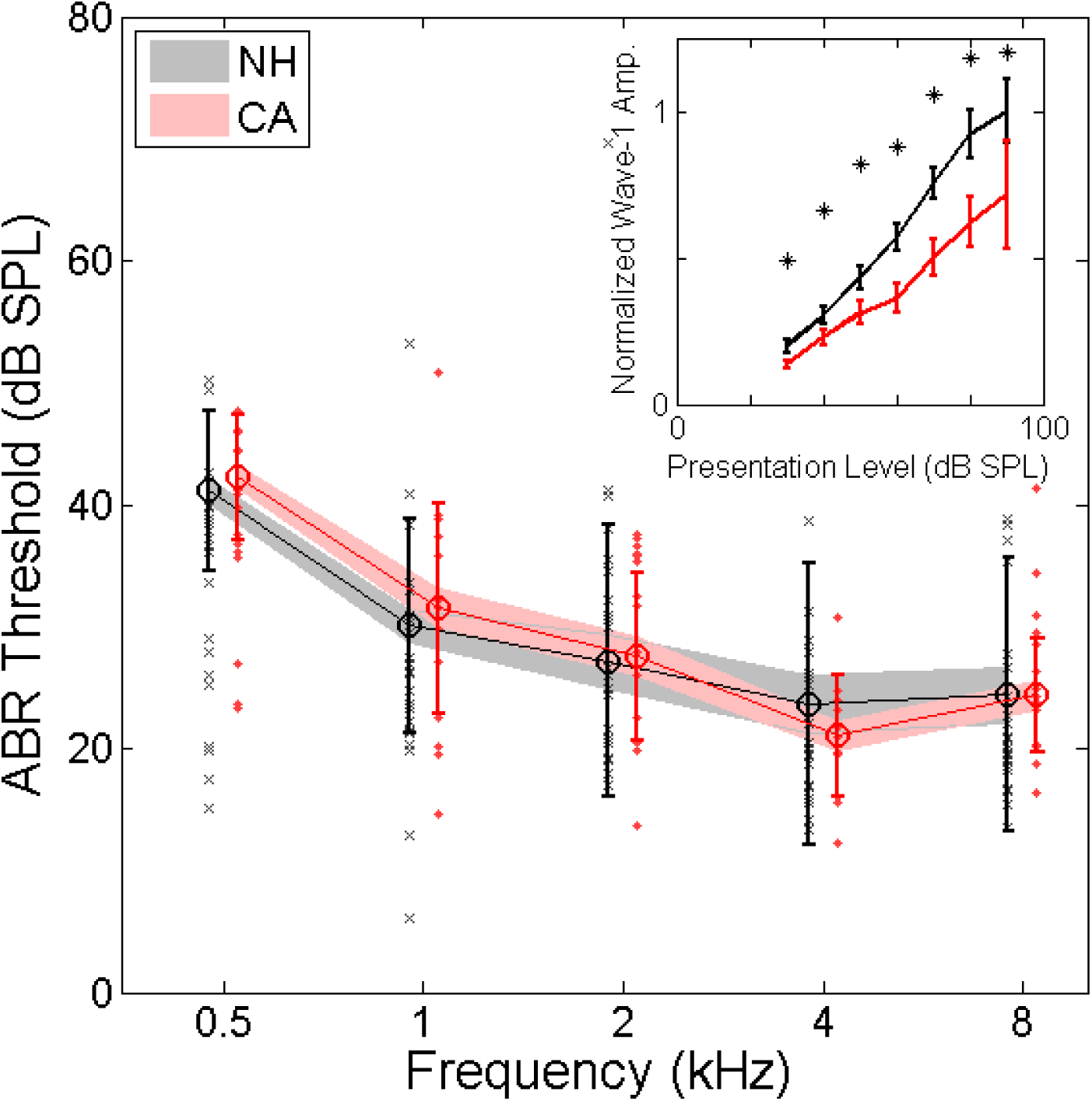
ABR thresholds plotted relative to tone frequency showed no significant difference between NH (black) and CA-exposed (red) animals. Solid lines and circles show mean threshold averaged across each group. Shaded regions depict 95% confidence intervals of the mean, and error bars show standard deviation. Grey x’s and red diamonds show amplitudes for individual NH- and CA-exposed animals, respectively. **Inset: Normalized wave-1 amplitudes were reduced following CA exposure.** Inset shows responses to a 4 kHz stimulus plotted against stimulus sound level and normalized relative to mean NH response at 90 dB SPL. At all sound levels, CA-exposed animals showed reduced wave-1 amplitudes.

### 3-3- Characterization of single-unit responses following carboplatin exposure

Average tuning-curve thresholds were slightly elevated (5-15 dB across CFs) but showed large variance (Fig. 6, A, Tukey’s HSD, *Q*=2.35, *P*<0.0001). Some CA-exposed fiber thresholds remained very sensitive (thresholds < 0 dB SPL) while others showed severe elevation (thresholds > 90 dB SPL). It’s assumed that low-threshold fibers are responsible for providing normal sensitivity to pure tones in quiet, as observed in ABR recordings, and that even a small number of fibers with “normal” thresholds is sufficient to preserve normal thresholds in quiet. Single-fiber tuning, measured as Q_10_ (= CF/BW_10dB_), was not significantly altered following exposure (Fig. 4, B, Tukey’s HSD, *Q*=2.35, *P*=0.18). Tuning curves of extremely impaired fibers still had the narrowly tuned “tip” around CF. This tip is the result of local mechanical action by OHCs. Previous studies using sound overexposure and aminoglycoside-induced ototoxicity have shown this tip is lost when OHCs are disrupted (Liberman, 1984; Liberman & Kiang, 1984). This further supports the argument that OHC function remains intact following the mild CA dose used here.

**Fig. 6.**
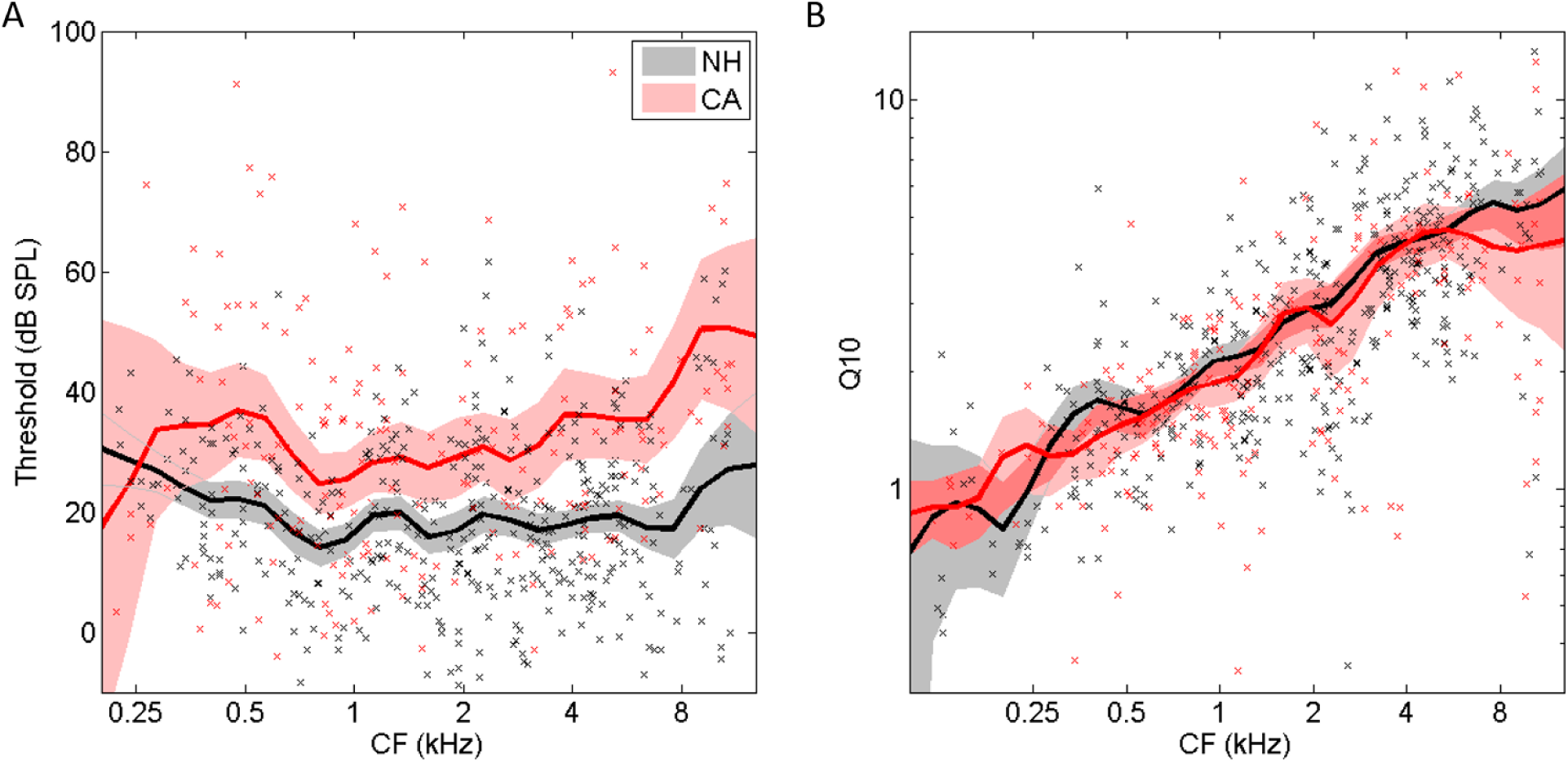
A) Average single-AN-fiber thresholds were slightly elevated following CA exposure. CA-exposed fibers showed a much wider range of thresholds compared to NH, with some fiber thresholds being as high as 90 dB SPL while others remained extremely sensitive. **B) AN fiber tuning was not altered following exposure as measured using the Q_10_ tuning sharpness metric.** Preservation of fiber tuning supports CA’s specificity towards IHCs. If OHCs were affected, this would likely be reflected in fiber tuning. X’s show individual fiber values for NH (black) and CA-exposed (red) fibers. Solid lines show octave-band averages for each group and shaded regions illustrate 95% confidence intervals of the mean.

On average across CA-exposed fibers, driven and spontaneous rates were reduced (Fig. 7). In NH ears, the distribution of SRs is typically bimodal with a peak at zero and another broader peak typically around 60 spikes/s. In CA-exposed ears, the distribution lacks the second peak at higher SRs and shows a stronger bias towards lower SRs. Across all fibers, mean SR in NHs was 40.9 spikes/s and 25.0 spikes/s in CA-exposed ears (Fig. 7, A). Using a commonly accepted SR-based categorization method (Liberman, 1978), we sorted fibers into low, medium, and high SR-fiber groups with a criterion of <.5 spikes/s, .5-18 spikes/s, and >18 spikes/s for each group, respectively. The NH population comprised 8% low-, 27% medium-, and 65% high-SR fibers, whereas the CA-exposed population comprised 16% low-, 35% medium-, and 49% high-SR fibers. The proportion of fibers in each SR group was shown to be significantly different following CA-exposure (*χ^2^*_2_ = 12.55, *P* = .0009). Mean saturated firing rates across all fibers in normal-hearing ears were 119.6 spikes/s, while in CA-exposed ears the mean saturated rate was 106 spikes/s (Fig.7, B, Student’s T-Test, *t_512_*=2.74, *P*=.0064).

**Fig. 7.**
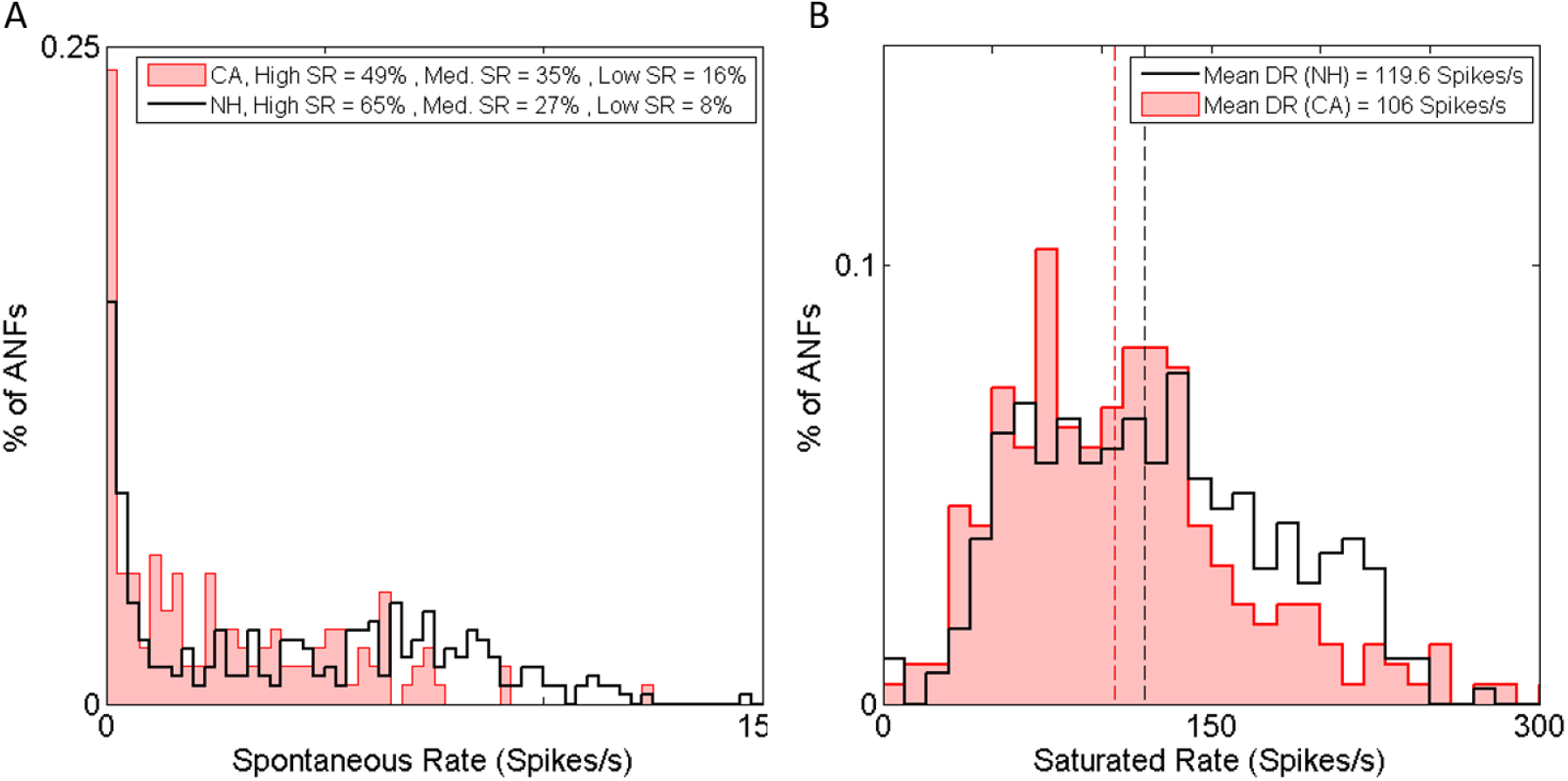
CA exposure reduced spontaneous firing rates (SR, A) and saturated firing rates (B). Histograms show distributions of firing rates for NH (black trace) and CA-exposed (red trace with pink fill) fibers. Vertical dotted lines (B) show the mean saturated rate for each group. Both SR and saturated rate distributions showed a significant shift towards lower rates following exposure.

Modulation synchrony to SAM tones in quiet, calculated as VS, showed no differences between NH and CA-exposed ears (Fig. 8, ANOVA, *F*_(1,3888)_=0.26, *P*=0.61). VS to both the carrier and modulator of SAM tones was unchanged following exposure. Strength of synchrony to the carrier was highly dependent on the CF of the fiber with strong phase locking observed in low-CF fibers and phase-locking beginning to roll off at a CF of 2 kHz (Temchin & Ruggero, 2010; Verschooten et al., 2019). At CFs above 2 kHz, phase-locking to the carrier is greatly reduced. Phase-locking to the 50-Hz modulator did not show any relationship to fiber CF, but was strongly dependent on presentation level, consistent with previous studies (Joris and Yin, 1992; Kale and Heinz, 2010). The strength of phase-locking to the modulator is non-monotonic, with synchronicity increasing at sound levels just above threshold to a maximal amount and then sharply rolling off at higher sound levels. The sound level that evokes maximum synchrony to the envelope is defined as the best modulation level (BML). In order to control for level-dependent effects, all subsequent SAM-tone stimuli were presented at the individual fiber’s BML. Average BML across fibers was not significantly different between NH and CA-exposed populations. DR at BML was significantly lower on average in CA-exposed fibers than NH, 36.4 spikes/s and 45.8 spikes/s, respectively (Student’s T-Test, *t_133_*=3.15, *P*=.002).

**Fig.8.**
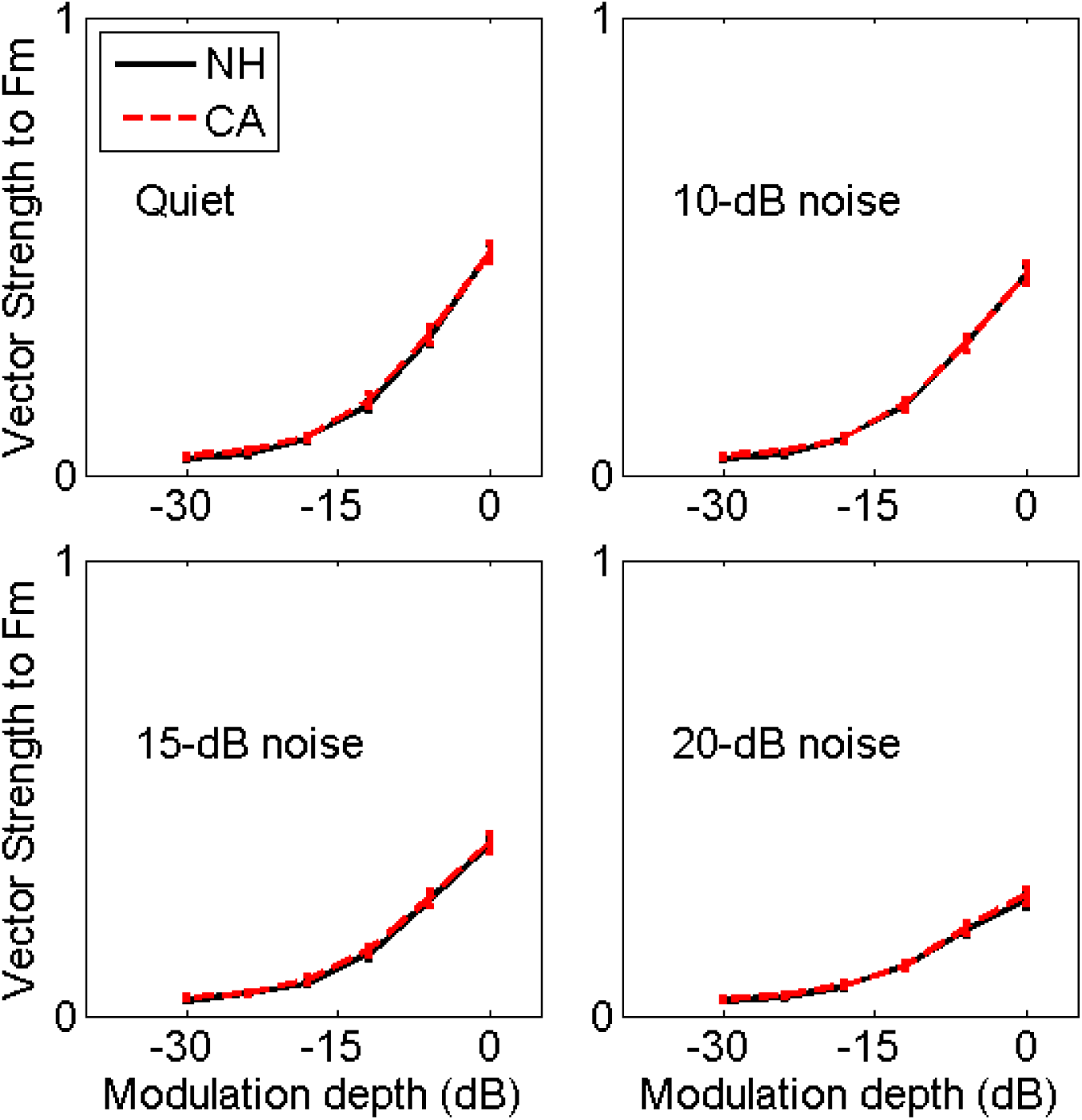
Average spike synchrony to the modulation frequency of SAM tones was not altered following CA exposure. Mean spike synchrony, measured as VS, was equally degraded in NH and CA-exposed fibers by decreasing modulation depth and increasing background noise. SAM tone stimuli were presented at the BML of each fiber.

### 3-4- Amplitude-modulation detection

Mean VS (and VSpp) to SAM tones of varying modulation depth were not different between NH and CA-exposed populations (Fig. 8). The presence of broadband noise (BBN) reduced synchrony to the envelope in both groups but there were still no differences between NH and CA-exposed fibers for all noise levels measured here. It is important to keep in mind that the VS metric reflects the mean synchrony averaged across many repetitions of the stimulus. In real-world environments, important stimuli are typically not repeated, and so stimulus information is redundantly encoded across multiple fibers (e.g., volley theory, (Wever & Bray, 1937)). When multiple fibers converge on individual neurons in the CN, the information of the fibers is integrated temporally. For reliable transfer of information, responses across fibers need to be consistent. Experimentally, this can be measured within individual fibers by monitoring the consistency of a single fiber’s response across multiple repetitions of the stimulus. Across CA-exposed fibers, mean VSpp did not change relative to NH fibers, but the variance of responses across repetitions was increased (Fig. 9).

**Fig. 9.**
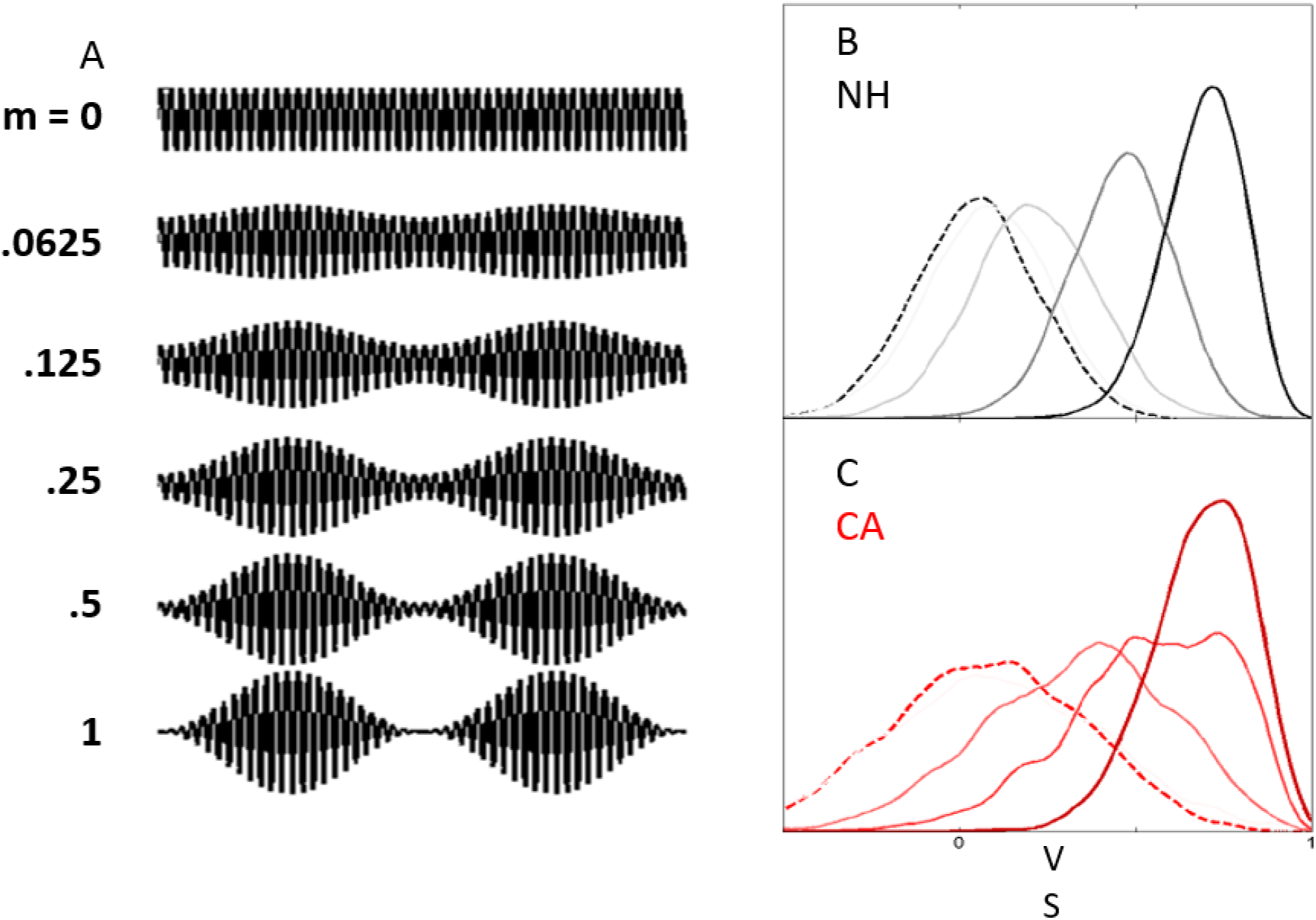
Average spike synchrony to amplitude modulation was not altered across NH and CA fibers, but reduced spikes led to increased variance across individual stimulus repetitions in CA-exposed fibers. A: Traces illustrating SAM stimuli of varying modulation depths, from completely unmodulated (top) to fully modulated (bottom). B and C: Example boot-strapped distributions of VSpp in response to modulated and unmodulated stimuli. Dashed lines indicate responses to an unmodulated tone. Solid lines indicate the response to modulated tones with increased line darkness indicating increased modulation depth. For each modulation depth shown, the CA distribution of VS was broader (higher variance) than the NH distribution.

Signal detection theory takes both means and variances into account when determining whether or not signals are discriminable from a given reference (in this case, the reference used for comparison was a pure tone at the fiber’s CF). Measuring how discriminable SAM-tone stimuli were from an unmodulated tone using the discriminability index (*d’,* Eq. 5) takes into account both the mean and the variance of a fiber’s responses across many repetitions and makes it possible to see what effect increased variance across repetitions due to reduced spike counts might have. When analyzed under this framework, differences in envelope coding between NH and CA-exposed fibers were revealed. Increased variance following CA exposure disrupts reliable coding of modulation-depth information and reduces fibers’ ability to detect modulation (Fig. 10). Statistical analyses help elucidate the source of these changes. In a simplified ANOVA model with drug exposure as the only factor, the effect of CA was highly significant (ANOVA, *F_1,3904_*=94.99, *P*<.0001). However, when DR was added as a factor, CA-exposure no longer had a significant effect on *d’* (ANOVA, *F_1,3888_*=2.95, *P*=.086) and DR was the main effect predicting discriminability (ANOVA, *F_1,3888_*=1606.14, *P*<.0001). This suggests that reduced DR following CA exposure leads to increased rep-to-rep variability, which reduces discriminability.

**Fig. 10.**
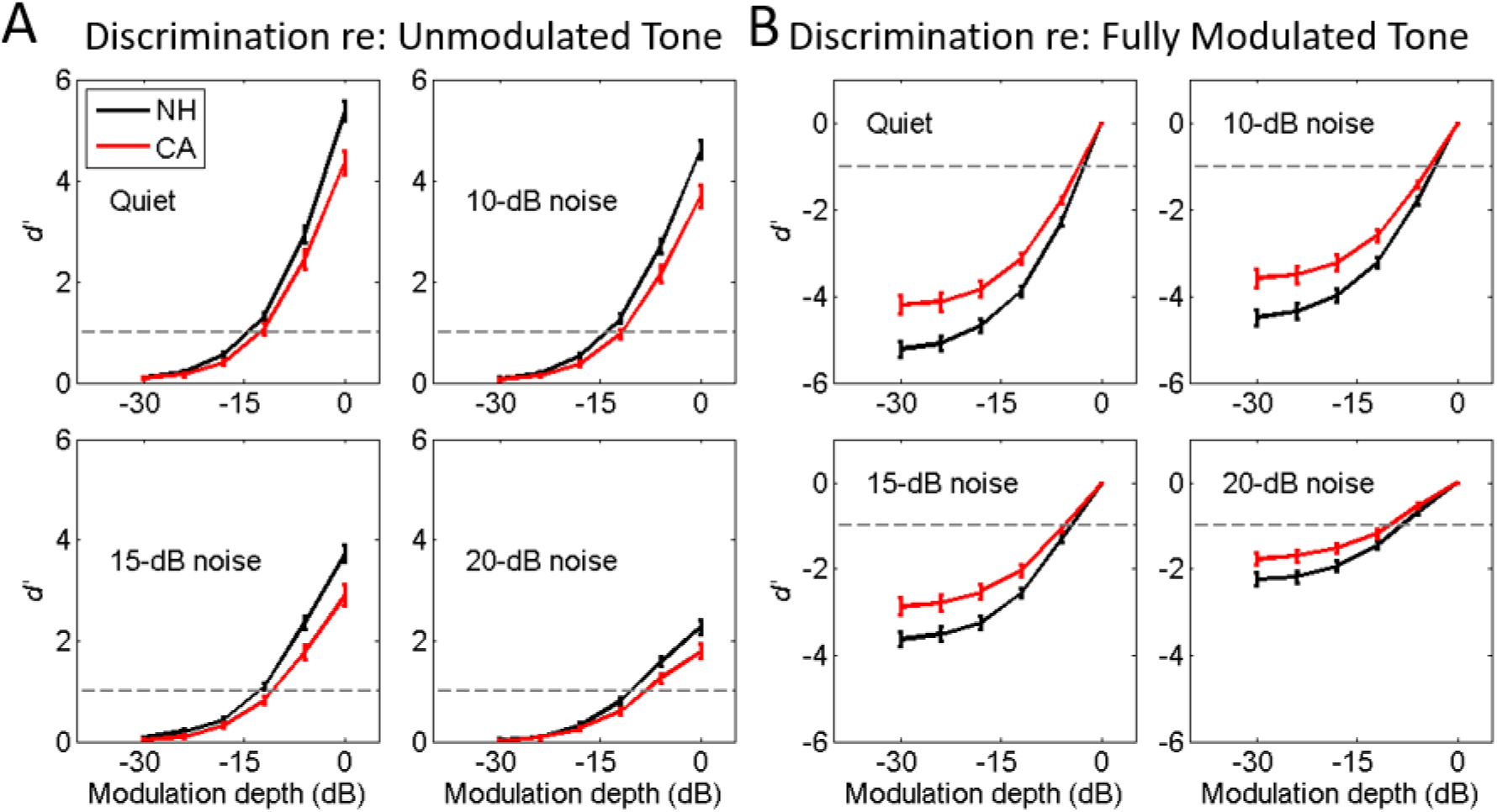
Detectability of amplitude modulation was degraded following CA exposure due to increased variability in CA-fiber responses that had lower spike counts. Detectability was calculated as the discriminability index (*d’* or “d-prime”) of a modulated-tone probe to an unmodulated-tone reference (A), or of a varyingly modulated tone probe to a fully modulated-tone reference (B, m=1). Each panel shows the neurometric functions in varying amounts of background noise. Discrimination index uses both the mean and the variability of responses to estimate signal salience. NH and CA fibers show equal mean synchrony to AM (see Fig. 8) so the observed reduction in *d*’ must be due to increased variability across boot-strap repetitions in CA-exposed fibers. The presence of background noise degraded the ability to discriminate for both NH and CA fibers but did not affect one group more relative to the other.

Average mutual information (MI) of VSpp and modulation depth was significantly reduced following exposure. Across NH fibers, average MI was .86 bits and across CA-exposed fibers average MI was .72 bits (Student’s T-Test, *t_133_*=3.33, *P*=.0011). The detectability of modulation, when compared relative to an unmodulated tone, was reduced in CA-exposed fibers. The threshold of modulation detection in quiet conditions for NH fibers was -11.3 dB and -9.8 dB in CA-exposed fibers, but this difference was not statistically significant (*T_104_* = 1.62, *P* = 0.1083). VS, d’, MI, and threshold of detection are all correlated with both SR and DR (See Table 1). VS is negatively correlated to both SR and DR. MI and d’ are negatively correlated to SR but positively correlated to DR (Table 1). Threshold of detection is positively correlated to SR and negatively correlated to DR (Table 1). In almost all cases, the strongest correlation is seen between these metrics and DR (Fig. 11, Table 1). This trend was present in both NH and CA-exposed fibers, which suggests that this is an intrinsic relationship in how ANFs encode AM stimuli. Since DR is the difference between the evoked firing rate during the stimulus period and the spontaneous rate during quiet, it can be interpreted as the relative number of spikes elicited by the stimulus above the “neural noise floor” of the spontaneous rate of the unit.

**Fig. 11.**
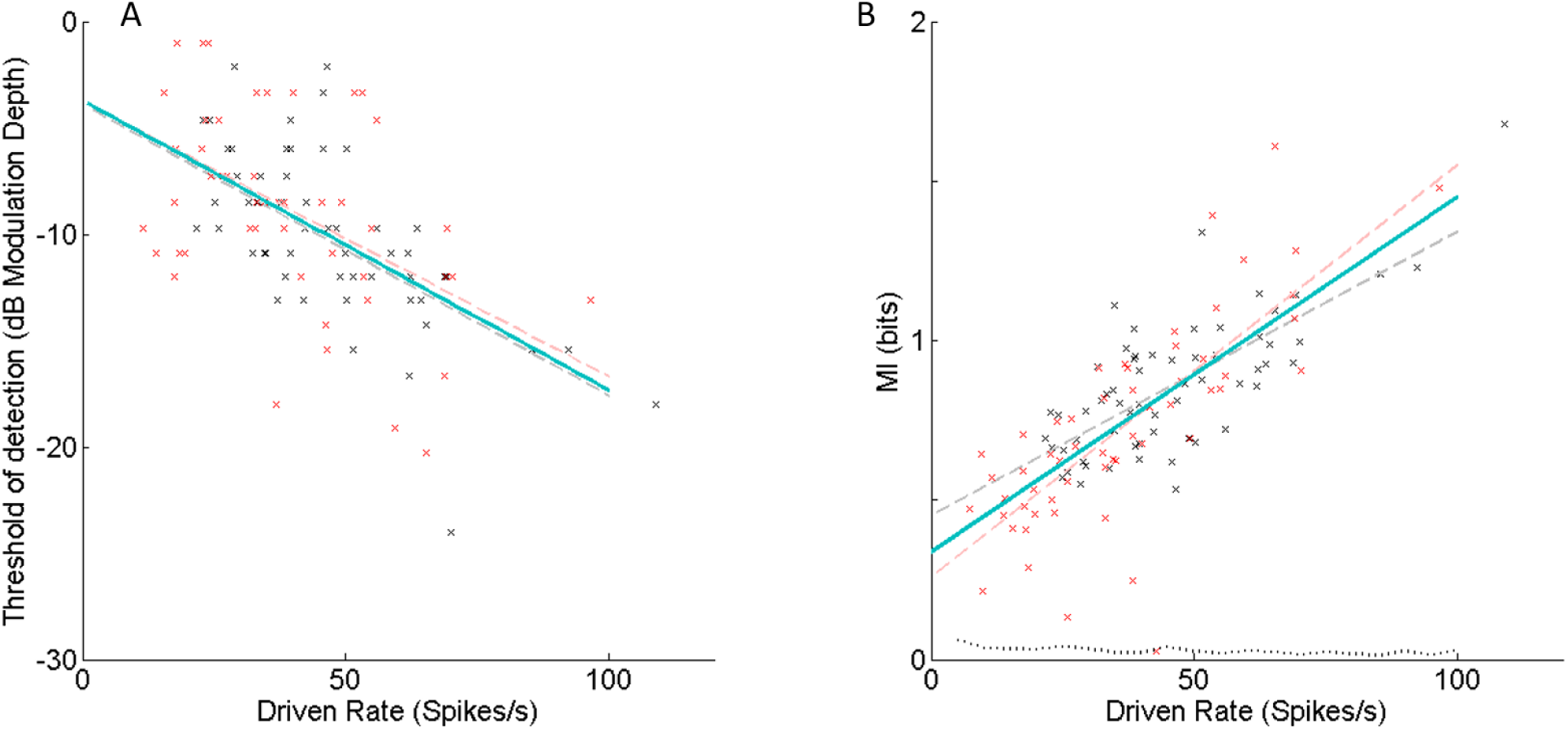
A fiber’s ability to efficiently encode amplitude modulation is strongly correlated to the DR of the fiber. A, threshold of AM detection. (the modulation depth where *d’*=1, statistically above chance). **B, mutual information between VSpp and modulation depth.** X’s show values for individual NH (black) and CA-exposed (red) fibers. Dashed lines show lines of best fit to each population, and the solid teal line shows the line of best fit across both populations. The dotted line in B shows the “Noise floor” of MI relative to DR calculated using randomly generated spikes uncorrelated to the stimulus. The strong correlation to DR was present in both NH- and CA-exposed fibers, which suggests that this is an intrinsic relationship in how ANFs encode AM stimuli. Furthermore, this would indicate that the reduced ability of CA-exposed fibers to accurately encode AM is due to their reduced output rather than an inherent difference in their temporal synchrony.

**Table 1.**
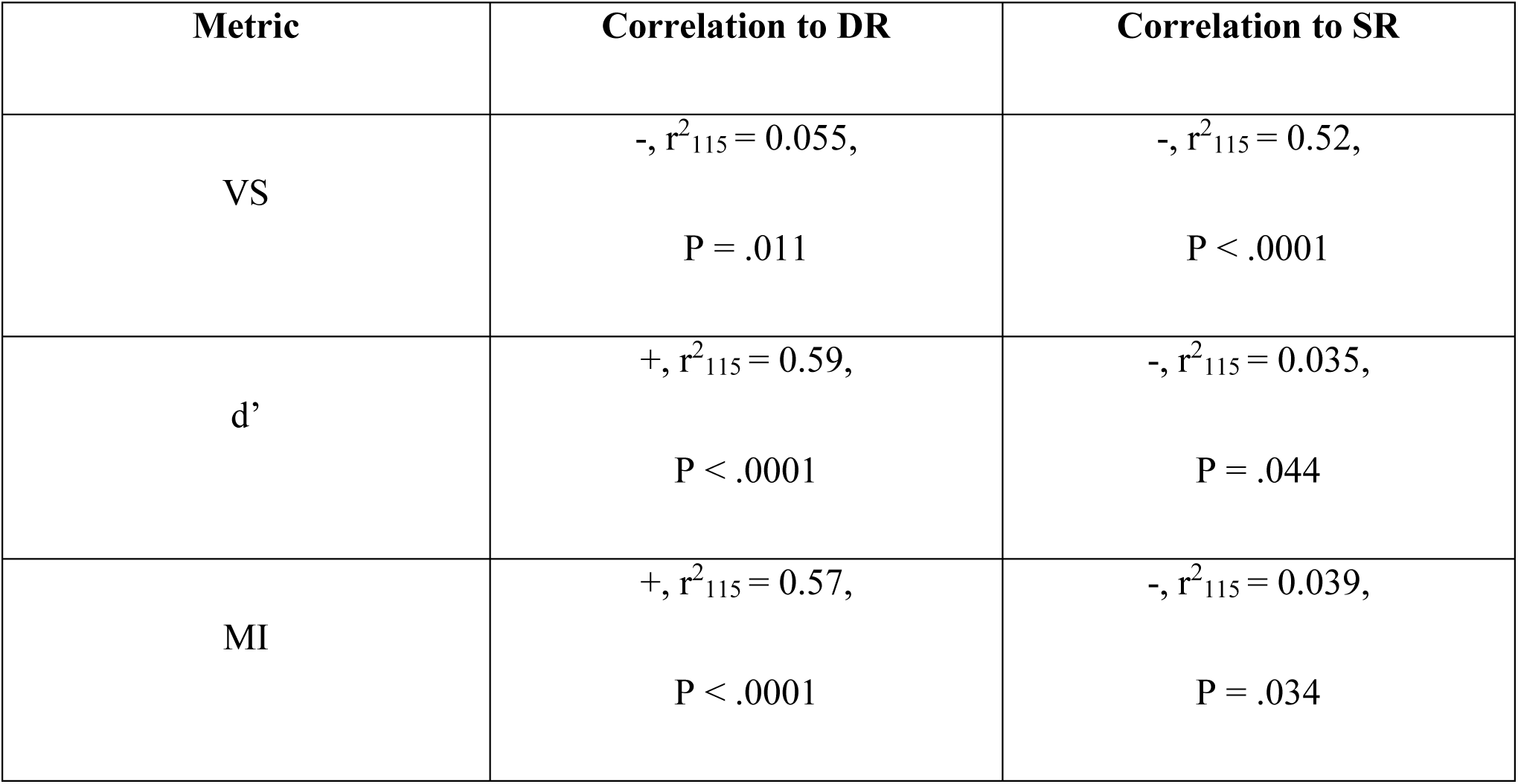
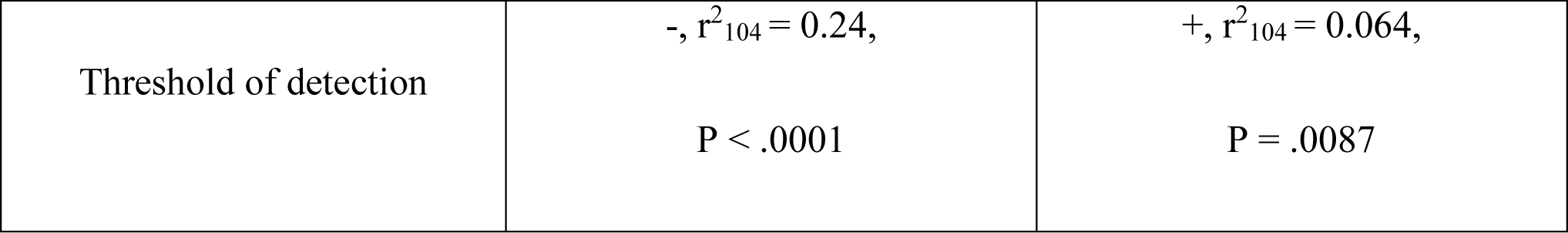
A summary of modulation-coding metric correlations with driven rate and spontaneous rate. + or – symbol indicates whether correlation was positive or negative. All correlations shown were significant.

The reason for reduced modulation depth detection capability in CA-exposed fiber is mathematically due to increased variability in modulation coding across repetitions. This increased variance arises because, following CA exposure, firing rates in exposed fibers are reduced. This causes an undersampling of the stimulus by the fiber and increases sampling bias in each repetition. This phenomenon was demonstrated by artificially limiting the number of spikes randomly selected during each bootstrap repetition for both NH and CA populations (Fig. 12). This essentially gives each fiber the same firing rate, no matter if it came from the NH or CA population. For this analysis, the number of spikes was set to 20 spikes per repetition or a DR of roughly 33 spikes/s. This amount corresponds with the lower end of DRs observed in CA-exposed fibers. When firing rate was controlled for in this manner, there was no distinguishable difference in the responses of the NH and CA-exposed fibers (Fig. 12). This observation suggests that the temporal response properties of the fiber have not changed following CA exposure, but rather by reducing the output of the IHC the relative amount of information that a given fiber receives is reduced and therefore its ability to reliably encode modulation depth information is impaired.

**Fig. 12.**
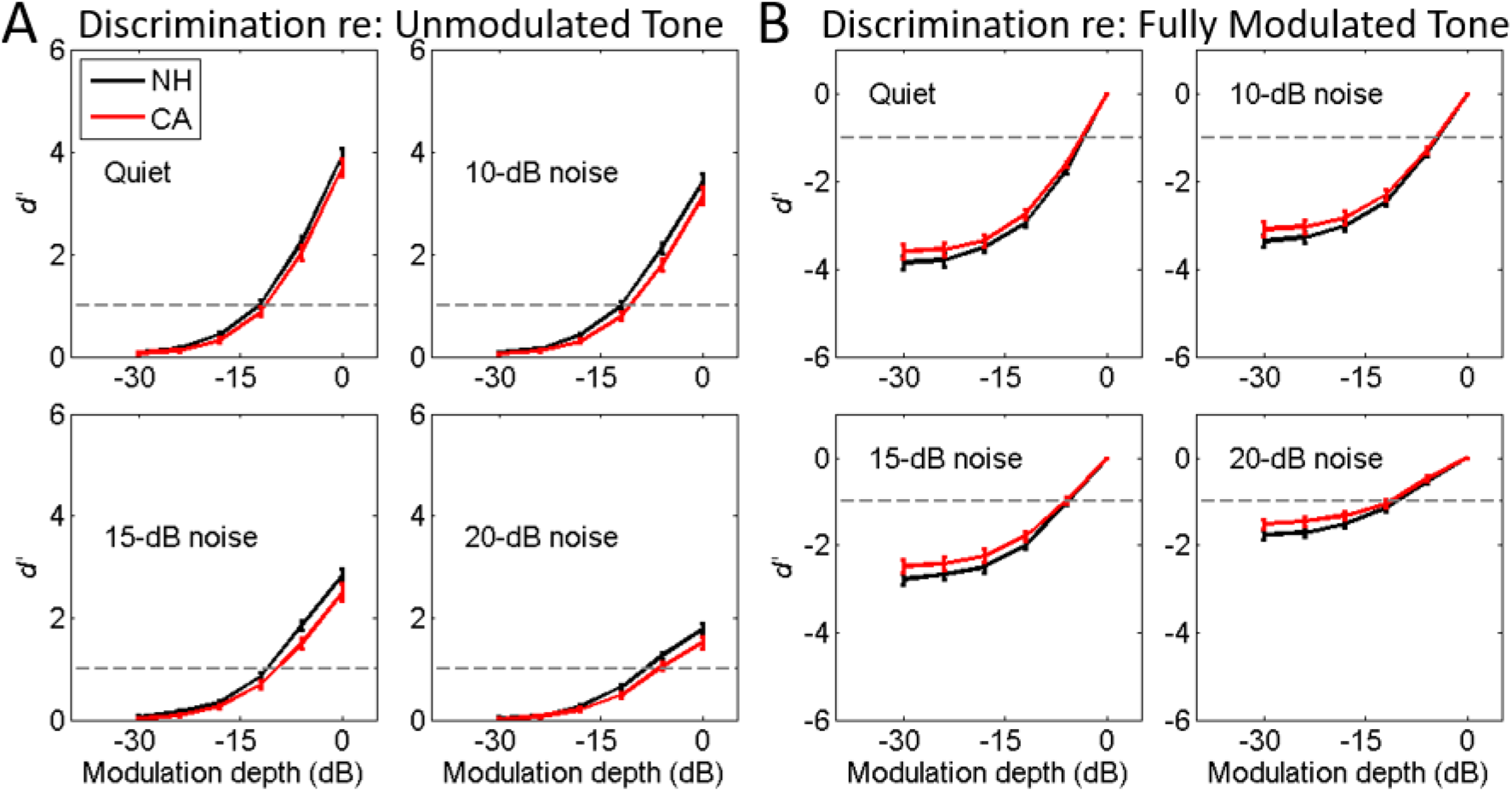
When spike count was controlled for, there was no difference in modulation detection ability between NH and CA fibers. When the number of spikes randomly picked for each boot-strapped repetition was held constant (n=20 spikes/rep) for all fibers in both NH and CA populations, there was no difference between the discrimination ability of NH and CA fibers. Left group shows comparisons between a pure-tone reference and probe tones of varying modulation, while left shows comparisons between a fully modulated tone reference and probe tones of varying modulation.

Across all the analyses within individual fibers discussed above, the presence of background BBN degraded accurate coding of AM in both groups but did not affect CA-exposed fibers any more strongly than NH fibers (ANOVA, F_1,3888_ = .0006, *P* = 0.98). This is likely due to narrow tuning of the BM being preserved following CA-exposure. This narrow tuning attenuates the majority of the energy carried by the noise.

### 3-5- Pooled-neurometric modulation detection

In order to investigate how reduced information at the level of individual AN fibers affects coding of modulation depth across fiber populations of varying size (Fig. 13), AN fibers were randomly selected into pooled populations and then put through a simulated two-choice modulation detection task. Population-pool sizes were set to 1, 2, 5, 10, 20, and 50 fibers per pool. In the case of pool size=1 (Fig. 13, A-D), essentially the capability of individual fibers to detect modulation, the capabilities of NH- and CA-exposed fibers closely followed the same trends observed in the d’-based tMDFs (see Fig. 10). CA-exposed fibers showed worse ability to detect modulation when compared to NH fibers (ANOVA, *F_2,25104_* = 46.57, *P* < .0001). Just as in the d’-based tMDFs, the presence of background BBN did not seem to affect CA-exposed fibers any more dramatically than it affected NH fibers (Fig. 10).

**Fig. 13.**
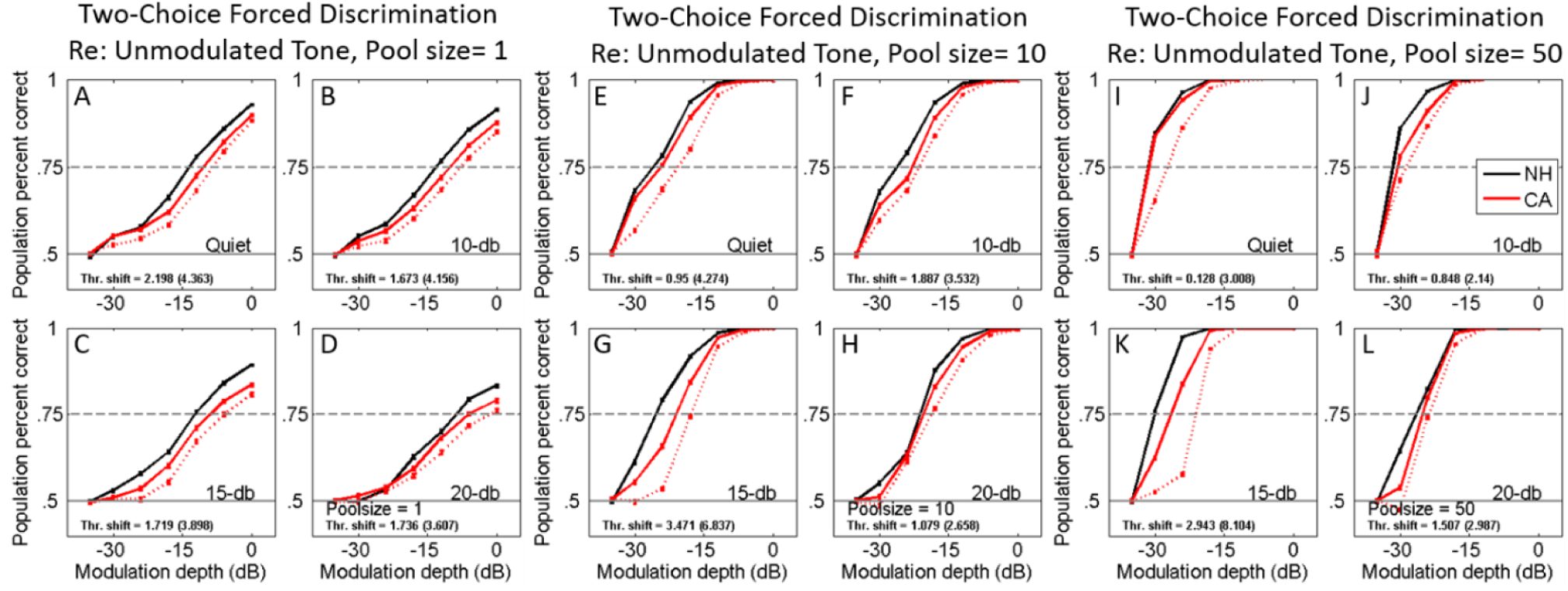
Pooled-neurometric analysis indicates that in quiet conditions a small number of “normal”-behaving fibers within the neuron pool are sufficient for normal AM detection, but when background noise is introduced and the simulated task becomes more difficult, the loss of redundancy causes reduced discrimination ability in pools of CA-exposed fibers. Pools were randomly chosen without replacement from three populations: NH (black lines), all CA-exposed fibers (solid red lines), and severely impaired CA-exposed fibers (dashed red lines, those with a DR of <40 Spikes/s). **A-D, in the case where pool size = 1, i.e., the performance of an individual fiber, the performance of the NH and all-CA groups was similar to that observed in the d’-based tMDFs (see Fig. 10)**. The presence of background noise impaired the performance of single-fiber pools for all groups but did not disproportionately affect any group. **E-L, as pool size is increased, performance improves for all groups, and with sufficiently large pool size (I-L), pools selected from across all CA-exposed fibers performed as well as NH in quiet (I).** However, when background noise was introduced the CA-exposed pools’ performance was more strongly degraded (J-L).

As the size of the pool of neurons was increased, the modulation depth capability of the neuron pools improved across all conditions. In the quiet condition at pool sizes above 20, CA-exposed pools performed just as well as NH pools (Fig. 13I). When background BBN was introduced (Fig. 13, J-L), the CA-exposed pools performed significantly worse than the NH pools. In the most severe BBN condition, 20 dB BBN re stimulus level (Fig. 13L), CA-exposed pools and NH pools both performed poorly with similar capabilities. This is likely some intrinsic floor effect of the noise.

A third category of pools was formed by sampling only those CA-exposed fibers with DRs < 40 spikes/s (approximately those with DRs below the mean of CA-exposed DRs). This category was formed to demonstrate the capabilities of the most impaired fibers exclusively (Fig. 13, dashed red line). In the case of pool size = 1, this category performed slightly worse on average than the full CA-exposed population, but the mean of the severely impaired group fell within the standard error of the full CA-exposed mean across all noise conditions. For higher pool sizes, the severely impaired group performed significantly worse than both NH pools and the pools formed from the full CA-exposed population (Tukey’s HSD, Q = 2.35, *P* < .0001). The differences between the full and severely-impaired CA-exposed populations indicate that it is possible for the more normally responsive fibers to “carry” the pooled response in quiet conditions. As the pool size increases, the chances of having more “good” fibers in that particular pool increase, and so the average success rate increases with pool size. In the pools of severely impaired CA fibers, there are no “good” fibers, and so across all conditions they perform more poorly than NH pools.

## 4- Discussion

### 4-1- IHC dysfunction is not reflected in evoked-potential thresholds but is observable in reduced ABR Wave-1 supra-threshold amplitudes sound levels

Preserved ABR thresholds following carboplatin exposure (Fig. 5) are consistent with other studies using carboplatin that have shown no significant elevation of pure-tone detection. In one such study, thresholds remained intact in behaving animals with as much as 80% IHC loss (Lobarinas et al., 2013, 2015). Reduced supra-threshold amplitude (Fig. 5 inset)is consistent with reduced neural output following CA exposure.

This reduction likely comes from two sources. The first is direct loss of IHCs. Based on the literature, the expected IHC loss is ∼20% for the carboplatin dosage used here (Trautwein et al., 1996). A representative cochleogram from our study showed ∼10-15 % IHC loss across the cochlea (Fig. 2). Without these IHCs the SGNs that innervated them will be without input and should no longer contribute to evoked population responses. The second source of reduced neural output likely comes from disruption of IHC stereocilia (Fig. 3), resulting in reduced ionic current through mechanically gated ion channels. Scanning electron microscopy showed disorganized stereocilia bundles following CA exposure (also see Wake et al., 1994). Dysfunction of mechanically gated ion channels would reduce acoustically driven currents, resulting in reduced synaptic output and consequently a reduced driven firing rate of innervating SGNs. Leak currents through mechanically gated ion channels are also one of several mechanisms behind spontaneous neurotransmitter release and spontaneous firing rates in SGNs. Similar to driven rates, reducing mechanically gated ion channels will result in reduced spontaneous firing in SGNs (Fig. 7A). It is important to note that this alteration in spontaneous firing is very likely mechanically distinct from the intrinsic synaptic properties that mediate the “type” of fiber (low, medium, or high SR fiber). By disrupting the stereocilia of the IHC, carboplatin exposure reduces current flow into the cell and, through this disruption, reduces synaptic output without necessarily altering the synapse itself.

It is quite likely that a patient who comes into the clinic presenting a phenotype similar to that observed in CA-exposed chinchillas would be diagnosed as normal hearing. If ABR thresholds are interpreted as being analogous to audiogram thresholds, then the average CA-exposed subject would present with normal audiogram thresholds and normal DPOAEs. The only indication of impairment that is directly measurable from the evoked potentials measured in this study is in the reduced amplitudes of ABR waves 1 and 2 at supra-threshold sound levels. This indicator is confounded by the lack of reduction in waves 4 and 5. A similar phenotype has been measured in the synaptopathic “hidden hearing loss” model (Kujawa & Liberman, 2009). Investigators of that model have hypothesized that following the loss of a significant portion of ANFs, central processing centers along the auditory pathway respond to the diminished input with increased central gain. This same hypothesis could be applied to the CA-exposed model of this study as well. Similar to the case of synaptopathy, input to the CNS from the auditory periphery has been reduced; therefore, if the hypothesis of increased centralized gain is true, it would almost certainly come into effect following CA-exposure. When interpreting the findings of the present study for the purpose of developing clinical assays of IHC-specific dysfunction, this presents an interesting problem. In humans, the dominant feature of the ABR waveform is wave-V. Wave I is typically small, and its amplitude can be difficult to reliably measure. A patient with a phenotype similar to the CA-exposed chinchillas would present normal wave-V amplitudes, and if wave I could not be reliably measured, then it is possible the clinician collecting the ABRs might dismiss reduced wave-I amplitude as measurement error. One method clinicians use to overcome the inherent patient-to-patient variability of ABR amplitudes is to use ABR-wave latency instead. These measures are much more repeatable and consistent across patients, but in the case of a CA-exposed subject, latency did not change (not shown), so this measure too would also fail to identify the dysfunction.

### 4-2- Degradation of amplitude-modulation detection stems from reduced spike rates

The lack of difference in threshold of modulation detection between NH and CA-exposed fibers is likely due to confounding influences of SR and DR. The detectability of modulation, d’, shows a negative correlation to SR, with lower SR fibers showing greater detection capability in both NH and CA-exposed fibers. Conversely, d’ shows a positive correlation to DR. Intuitively, these relationships makes sense. Spontaneous spikes are events uncorrelated with an input stimulus and thus do not convey useful information to processing centers higher in the auditory pathway. Experimentally, SR is measured by recording activity in the absence of stimulation, but the leak currents that cause these spontaneous events are still present during stimulation. It is likely that these currents result in spikes uncorrelated to the stimulus, contributing noise to the response and weakening the strength of synchrony. Fibers with lower spontaneous rates have less of this “neural noise,” and as such, the information carried by each spike is more reliable and more strongly correlated to the stimulus. In order to account for the presence of these uncorrelated spikes when calculating DR, SR is subtracted from the measured firing rate during the stimulus period. Because of this, the relationship between DR and modulation coding is more complicated. Average synchrony to the modulator, VS, is lower in fibers with higher DRs, similar to SR, but detectability of modulation, d’, which is derived from VS, is greater in fibers with higher DRs. The reason for these seemingly contradictory results is that these two metrics measure different things. VS measures the precision with which spikes synchronize to a periodic component of the stimulus. To achieve maximum VS of 1 would require no more than 1 spike per cycle to be fired. This almost never occurs in a fiber’s responses to SAM tones because the fiber is responding to both the modulator and to the carrier components. At low modulation frequencies, a single cycle of the stimulus is long enough that multiple spikes are fired during its period. Having these extra spikes, even though they are correlated to the modulator, limits the calculated VS. These extra spikes, however, can be important for discriminating between different modulation depths. A larger number of stimulus-evoked spikes provides more observations of the stimulus-related temporal response, allowing differences between modulation depths to be estimated more reliably. Conversely, the reduced driven rates of CA-exposed fibers provide fewer stimulus-evoked spikes on which to base this discrimination, thereby reducing neurometric modulation sensitivity. This property is demonstrable using physiological data by artificially limiting the number of spikes per bootstrap repetition with both NH and CA-exposed data so that the number of driven spikes per bootstrap repetition is the same in both cases (Fig. 10). When the differences in firing rates are controlled in this manner, then the responses of NH and CA-exposed fibers are indistinguishable from one another.

### 4-3- Pooled-neurometric modulation detection suggests that redundancy of information is critical for accurate modulation depth coding in noisy conditions

The auditory periphery has a large amount of built-in redundancy. Each IHC is innervated by ∼15-30 ANFs. The basilar membrane possesses amazing frequency selectivity, but even for a relatively quiet pure tone, multiple nearby IHCs will be activated, and thus as sound level increases, excitation will spread along the basilar membrane. As electrically active cells, neurons and hair-cells have very high metabolic demands compared to other varieties of cells. It stands to reason that if the expensive energy demands of the redundancy built into the auditory periphery have been conserved throughout the history of mammalian evolution, then that redundancy is beneficial. Using pooled-neurometric analysis, we can demonstrate the importance of redundancy in the task of modulation detection in two ways. First, by increasing the number of fibers in the pool, performance improves across all cases. This makes sense intuitively because each additional fiber added to the pool can be thought of as an additional chance to “hear” the stimulus. Statistically, this makes sense within the framework of the metric itself. Neuronal pools are randomly sampled from the total population. The more fibers sampled within the pool, the closer the mean response of the pool will likely be to the mean response of the population. In essence, a larger pool size is analogous to a larger sample size during an experiment and increases the reliability of the findings.

The second manner in which redundancy’s importance is demonstrated is by the difference in performance between the NH and CA-exposed populations and between the full CA-exposed population and the more severely impaired subset. In the case of individual fibers (pool size=1), NH outperforms CA-exposed fibers. However, as the pool size increases, the performance of both groups improves, but the performance of CA-exposed pools becomes closer and closer to that of NH pools until, at higher pool sizes (≥20), the performance of both groups is practically identical. Conversely, in the single fiber pool case, the severely impaired subset of CA-exposed fibers performs more poorly than the full CA-exposed population, but the two populations still fall within the standard error of each other. As the pool sizes increase, the full population’s performance moves closer to that of the NH, but the severely impaired subset’s performance remains worse at all pool sizes and conditions. The differences at higher pool sizes in the percent-correct scores of pools from the full CA-exposed population and those formed from the most severely impaired CA-exposed suggest that as pool size increases, the best-performing fibers in the pool are able to “cover” for the more poorly performing fibers present in the pool. In a large pool of fibers, there will be a sufficient amount of “normal-behaving” CA-exposed fibers that they are able to perform as well as NH pools in quiet conditions. In noisy conditions, this redundancy is especially important. The presence of background noise degrades mean spike synchrony (VS_pp_) equally in NH and CA-exposed fibers (Fig. 6). Similarly, when background noise is present, the neurometric performances of both NH and CA-exposed individual fibers are degraded proportionally to the same degree by the noise (Fig. 8 and Fig. 11, A-D). In the pooled-neurometric paradigm, adding additional fibers improves percent correct scores substantially (Fig. 11, E-L) for both NH and CA-exposed fiber-pools in quiet conditions, but CA-exposed pools benefit less from higher pool size in noisy conditions. The addition of background noise makes the simulated task more difficult, and the small number of “normal behaving” CA-exposed fibers within the CA-exposed pools is not sufficient to match the performance of NH pools. When pools are made up of only severely impaired CA-exposed fibers, there are no “normal behaving” fibers, and so this group consistently performs more poorly than the other two groups even in the quiet condition.

## Acknowledgment

This work was supported by NIH NIDCD grants T32-DC00030.(D,A), R01-DC009838 (M.G.H.), and F32-DC022782 (A.F.). Electron microscopy and SEM sample preparation were performed using instrumentation in the Purdue Electron Microscopy Center (RRID:SCR_022687).

